# A close unicellular relative reveals aggregative multicellularity was key to the evolution of animals

**DOI:** 10.1101/2025.05.14.654023

**Authors:** Ruibao Li, Jennah E. Dharamshi, Kyle Kwok, Iñaki Ruiz-Trillo, Joseph P. Gerdt

## Abstract

How animals evolved complex multicellularity from their unicellular ancestors remains unanswered. Unicellular relatives of animals exhibit simple multicellularity through clonal division, formation of multinucleate coenocytes, or aggregation.^1^ Therefore, animal multicellularity may have evolved from one (or a combination) of these behaviours. Aggregation has classically been dismissed as a means to complex multicellularity.^2^ However, aggregation occurs in many extant animal cells and has also been recently described in three different unicellular relatives of animals (the choanoflagellates *Salpingoeca rosetta* and *Choanoeca flexa*, and the filasterean *Capsaspora owczarzaki*).^3–6^ It is unclear whether aggregation in these species is derived or ancestral, and its relevance for animal origins remains unknown. To fill this gap, we investigated whether an additional unicellular relative of animals can undergo aggregation. We discovered that the marine free-living bacterivorous filasterean *Ministeria vibrans*^7^ forms homogeneous aggregates with reproducible kinetics that have long-term stability when cultured with an alphaproteobacterium. We found that many multicellularity genes involved in animal cell adhesion, signaling, and transcriptional regulation were deployed during this process. Our findings suggest that the last unicellular ancestor of animals had the capacity to aggregate using key animal multicellularity genes and that improved feeding and sexual reproduction may be evolutionary drivers of this aggregation.

## Introduction

How animals (Metazoa) evolved from their unicellular ancestors remains a mystery. One of the most contentious debates about the transition is how the last unicellular ancestor of animals looked and behaved. Data from diverse unicellular animal relatives (i.e., choanoflagellates, filastereans, ichthyosporeans, and corallochytreans, which together with Metazoa form the Holozoa clade, Fig. 1a) suggest that the last unicellular animal ancestor had the capacity for simple regulated multicellularity. Namely, several choanoflagellate species can form multicellular colonies through clonal division, the filasterean *Capsaspora owczarzaki* can form temporary multicellular aggregates, and most ichthyosporeans and corallochytreans can form multinucleated coenocytes.^5,6,8–21^

**Figure 1.**
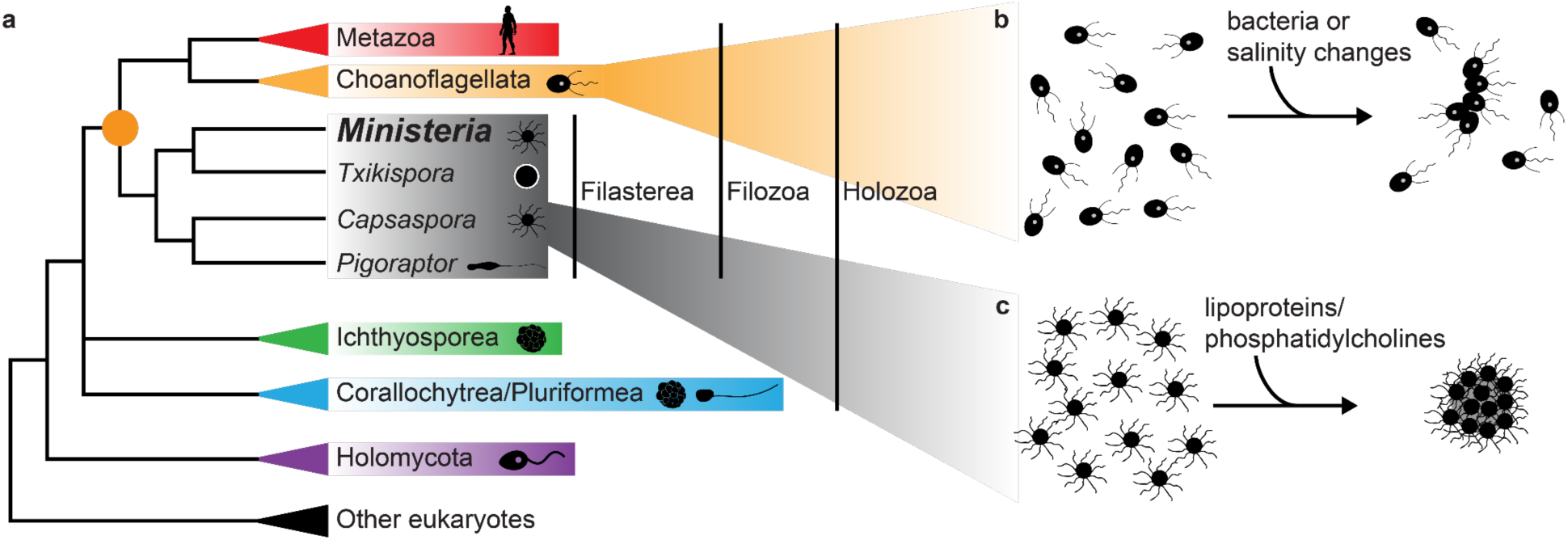
Multicellular aggregation in unicellular holozoans. a) Schematic phylogeny of holozoans. Orange circle represents the ancestor of Filozoa–the clade that comprises metazoans, choanoflagellates, and filastereans. b) The choanoflagellate *S. rosetta* forms moderately sized, stringy, multicellular aggregates in the presence of bacterial chondroitinases.^3^ A fraction of *C. flexa* choanoflagellate cells also aggregate upon changes in salinity.^4^ c) The filasterean *C. owczarzaki* homogeneously forms large multicellular aggregates in the presence of lipoproteins and phosphatidylcholine lipids from animal hosts.^5,6^ Neither aggregative process has been monitored transcriptionally or proteomically across the entire range of development to determine which animal multicellularity genes are essential for the initial formation and maturation of aggregates in filozoans.

Aggregation has been largely neglected as a foundation for evolving multicellularity in the first animal. Evolving complex multicellularity through aggregation has often been excluded due to the theoretical costs of preventing cheating.^22^ However, finding multicellularity through regulated cell aggregation in a close relative of animals,^4–6,11^ together with research showing that spatiality combined with group selection facilitates the survival of cooperators against cheaters in an aggregate, challenged those views.^23^ However, thus far our knowledge on aggregation in unicellular animal relatives is limited and primarily based on the filasterean *C. owczarzaki* and two case studies in choanoflagellates.^3–6,11^ Since *C. owczarzaki* aggregates in response to lipid cues from its putative animal host,^5^ this behavior may have been a relatively recent adaptation to its lifestyle as a symbiont. The choanoflagellate *Salpingoeca rosetta* can form small aggregates in response to *Vibrio* bacteria^3^ and *Choanoeca flexa* can form moderately sized aggregates in response to a drop in salinity,^4^ but these behaviours have not been studied extensively (Fig. 1b–c). Furthermore, gene expression has not been studied dynamically throughout the process of aggregation in any of these three organisms. Therefore, it is still unclear if aggregation was ancestral in unicellular animal relatives and whether genes critical for animal multicellularity and development are deployed during their aggregative behaviors. If they are, it would suggest that the unicellular ancestor of animals had the capacity for multicellular aggregation.

To address this question, here we studied *Ministeria vibrans*, a distant filasterean relative of *C. owczarzaki*.^24,25^ *M. vibrans* is a marine free-living bacteriovore—the lifestyle that is typically ascribed to the last unicellular ancestor of animals.^26,27^ Finding aggregation in a second filasterean (which is free-living) would suggest that aggregation is ancestral in Filozoa (Fig. 1a). In the original description of *M. vibrans*, occasional tiny cell clumps were observed (∼5 cells per clump). However, no remarkable multicellular aggregates were described.^7,28^ To determine if aggregation is an ancient behavior conserved in Holozoa, we asked if *M. vibrans* also has the capacity to form multicellular aggregates, and if so, which genes were regulating such behaviour. We found that *M. vibrans* formed large aggregates that persisted for over 60 days when cultured with a specific alphaproteobacterium, *Thalassospira lucentensis*. The entire culture homogeneously aggregated with consistent kinetics. We assessed gene expression profiles at six different time points during the *M. vibrans* aggregation process and found that many genes associated with animal multicellular cell adhesion, signalling, and transcriptional regulation were upregulated. The finding suggests that the last unicellular ancestor of animals had the capacity to form multicellular aggregates, and that to do so, it used genes crucial for multicellularity and development in extant animals.

## Results

### *Ministeria vibrans* multicellular growth is triggered by cultivation with a bacterium

To leverage *M. vibrans* as a model for the involvement of cellular aggregation in a free-living unicellular holozoan, we first needed to find a condition in which it reproducibly aggregated. In line with other reports of only occasional tiny clusters of a few cells,^28,29^ we too found no significant aggregates in the *M. vibrans* culture obtained from the American Type Culture Collection (ATCC #50519). Given the previous finding that *C. owczarzaki* aggregation is induced by lipoprotein complexes,^5,6^ we applied low-density lipoproteins (LDL) to the *M. vibrans* culture. However, this treatment failed to induce aggregation (Extended Data Fig. 1).

Since LDLs did not induce *M. vibrans* aggregation, we next considered that *M. vibrans* may aggregate in response to the presence of specific bacteria, as observed for the choanoflagellate *Salpingoeca rosetta*.^3^ We first assessed which bacteria were present in the original *M. vibrans* culture using 16S rDNA amplicon sequencing and found 45 unique bacteria, with *Thalassospira* being the most abundant (63%, Extended Data Fig. 2 and Data S1). We isolated colonies of a single *Thalassospira* bacterium (*T. lucentensis* Mv1, Data S2) from the *M. vibrans* culture and re-introduced it into flow-cytometry-purified *M. vibrans* to generate a monoxenic culture (Extended Data Fig. 2). The monoxenic culture was verified by 16S rDNA amplicon sequencing, which revealed no other bacteria present (Extended Data Fig. 2 and Data S3). The monoxenic culture grew well—in fact better than the original xenic culture (Fig. 2a). Remarkably, in the monoxenic culture, *M. vibrans* formed large multicellular clusters (Fig. 2b–c) that resembled those formed by *C. owczarzaki.*^5,6^ Therefore, monoxenic cultivation with the alphaproteobacterium *T. lucentensis* triggers multicellular growth in *M. vibrans*.

**Figure 2.**
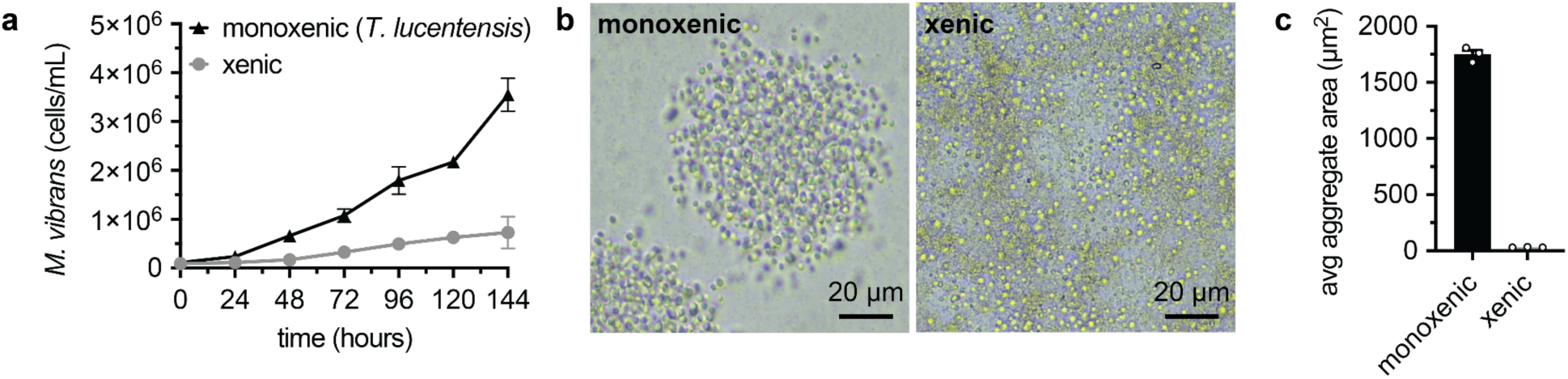
*M. vibrans* forms multicellular clusters in monoxenic culture with an alphaproteobacterium. a) Growth curves of the monoxenic *M. vibrans* culture with only *T. lucentensis* bacteria compared to the original xenic culture. Scale bars represent standard error of the mean (SE) of a biological triplicate. b) Representative images of aggregates formed in the monoxenic culture compared to the original xenic culture. c) Average aggregate area quantified for the monoxenic *M. vibrans* culture compared to the original xenic culture. Scale bars represent SE of a biological triplicate. Individual replicates are displayed with circles.

### *Ministeria vibrans* cells homogeneously aggregate with reproducible kinetics

To assess the potential of *M. vibrans* for investigating the involvement of multicellular genes in holozoan cellular aggregation, we needed to confirm that its multicellular clusters were formed through aggregation. We first monitored the process of cluster formation over time, which revealed that small clusters of dozens of cells merged into larger multicellular clusters containing hundreds of cells (Fig. 3a–b and Video S1). The observation matched previous observations in *C. owczarzaki*,^5,6^ suggesting that the large cell clusters were formed by aggregation. We also noted that the timing of cluster formation appeared too fast to arise from a single cell undergoing multiple rounds of cell division. Clusters containing dozens of cells formed within 24 h, which is only enough time for one or two rounds of cell division for *M. vibrans* (Fig. 2a and 3b). Furthermore, we observed that aggregates could form even when replication was blocked by aphidicolin (Fig. 3c, Extended Data Fig. 3, and Video S2), showing that daughter cell formation is not required for cell cluster formation. Together, these experiments demonstrate that *M. vibrans* forms large clusters of hundreds of cells through aggregation.

**Figure 3.**
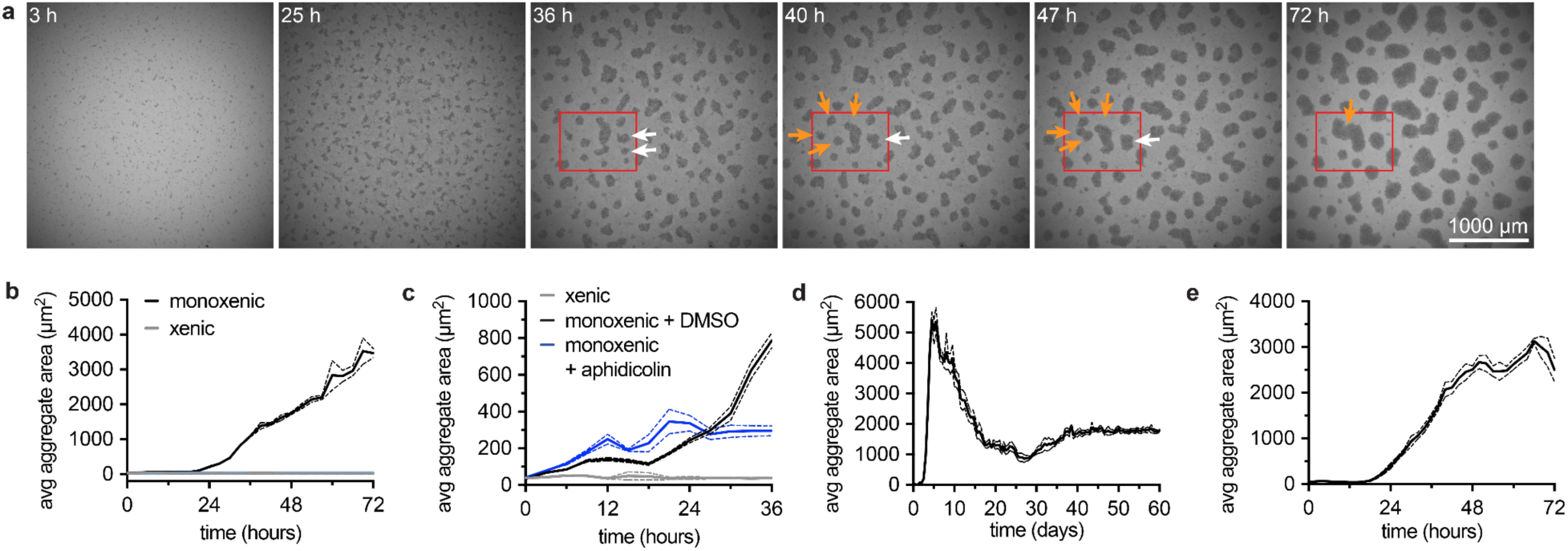
*M. vibrans* forms homogenous multicellular clusters through aggregation. a) Selected images of the formation of multicellular aggregates by the monoxenic *M. vibrans* culture after dilution into fresh growth media (see Video S1). Coloured arrows in the red box highlight small aggregates merging into large aggregates. b) Average aggregate area quantified for *M. vibrans* cultures (monoxenic vs xenic control) over time after dilution into fresh growth media (see Video S1). Dashed lines represent standard error of the mean (SE) of a biological triplicate. c) Average aggregate area quantified for the *M. vibrans* cultures incubated with aphidicolin (an inhibitor of replication, see Extended Data Fig. 3 and Video S2). Controls are also shown for non-treated cultures (monoxenic and xenic). Dashed lines represent SE of a biological triplicate. d) Average aggregate area quantified for the *M. vibrans* monoxenic culture over 60 days (1,440 h) after dilution into fresh growth media (see Video S3). Dashed lines represent SE of a biological triplicate. e) Average aggregate area quantified for the *M. vibrans* DSM 114853 strain in monoxenic culture with *T. lucentensis*, monitored for 3 days (72 h, see Video S4). Dashed lines represent SE of a biological triplicate.

In the related filasterean *C. owczarzaki*, aggregates appear to form through filopodial contacts that contract, pulling the individual cells into tight clusters.^6,30–32^ Likewise, analysis of scanning electron microscopy (SEM) images of *M. vibrans–T. lucentensis* cultures showed filopodia of different lengths connecting adjacent cells within aggregates (Extended Data Fig. 4a). The filopodia appeared to wrap around each other, generating extended surface areas of contact along each filopodium (Extended Data Fig. 4a). In many cases, the entangled filopodia appeared to have retracted almost completely (Extended Data Fig. 4b), forming very tight interactions between cells—as seen in *C. owczarzaki* aggregates.^11^ Intriguingly, *M. vibrans* aggregates in the *T. lucentensis* monoxenic culture persisted for at least 60 days (Fig. 3d and Video S3), which contrasts with *C. owczarzaki* aggregates, which disperse after 1–3 days—presumably upon depletion of the aggregate-inducing lipids.^5,6,33^ Perhaps a yet-unknown inducer of *M. vibrans* aggregation is continuously produced by *T. lucentensis*, affording persistent aggregation.

To examine if aggregative behavior is universal among *M. vibrans*, we acquired a second *M. vibrans* isolate (*M. vibrans* DSM 114853) and generated a monoxenic culture with *T. lucentensis* (Data S4). It aggregated in a very similar fashion (Fig. 3e). Therefore, *M. vibrans* consistently forms large multicellular aggregates when cultured with the alphaproteobacterium *T. lucentensis*.

### The dynamics of *M. vibrans* aggregative multicellularity

For several reasons, the *M. vibrans* aggregative phenotype is ideal for assessing whether cell aggregation could have played a role in animal origins. First, *M. vibrans* is a close unicellular relative of animals, distant from but related to *C. owczarzaki*, sharing several “multicellular” genes with animals.^18,29,34^ Second, it has a free-living bacterivorous lifestyle that may match the lifestyle of the last unicellular animal ancestor. Finally, the monoxenic culture homogeneously aggregates with reproducible kinetics and is stable over longer time periods, allowing us to sequence population-level transcriptomes at defined times to assess gene expression of *M. vibrans* as single cells, small initial aggregates, and large mature aggregates.

To identify genes involved in and dynamically expressed during the process of *M. vibrans* aggregation, we performed a time series experiment with 24 monoxenic cultures of *M. vibrans* growing with *T. lucentensis*. RNA was extracted and sequenced from four replicate cultures each at different stages along the formation and maturation of aggregates: 12 h, 18 h, 24 h, 33 h, 48 h, and 72 h after synchronous inoculation in fresh media (Fig. 4a, Extended Data Fig. 5a, and Data S5-7). Aggregates began to form after approximately 18 h in all replicates and continued to grow in size throughout the remainder of the time series, with aggregates becoming more stable in size from approximately 48 h. Validating our experimental approach, analyses of sample relationships revealed that the gene expression profiles from replicate time point cultures were most similar to each other, with the greatest variation at 18 h, just prior to the initiation of aggregation (Fig. 4b and Extended Data Fig. 5b). Replicates taken closer together in time have more similar gene expression profiles, with the largest shifts in gene expression occurring between 12 h and 24 h during the initiation of the aggregation process, and between 33 h and 48 h during the establishment of large mature aggregates.

**Figure 4.**
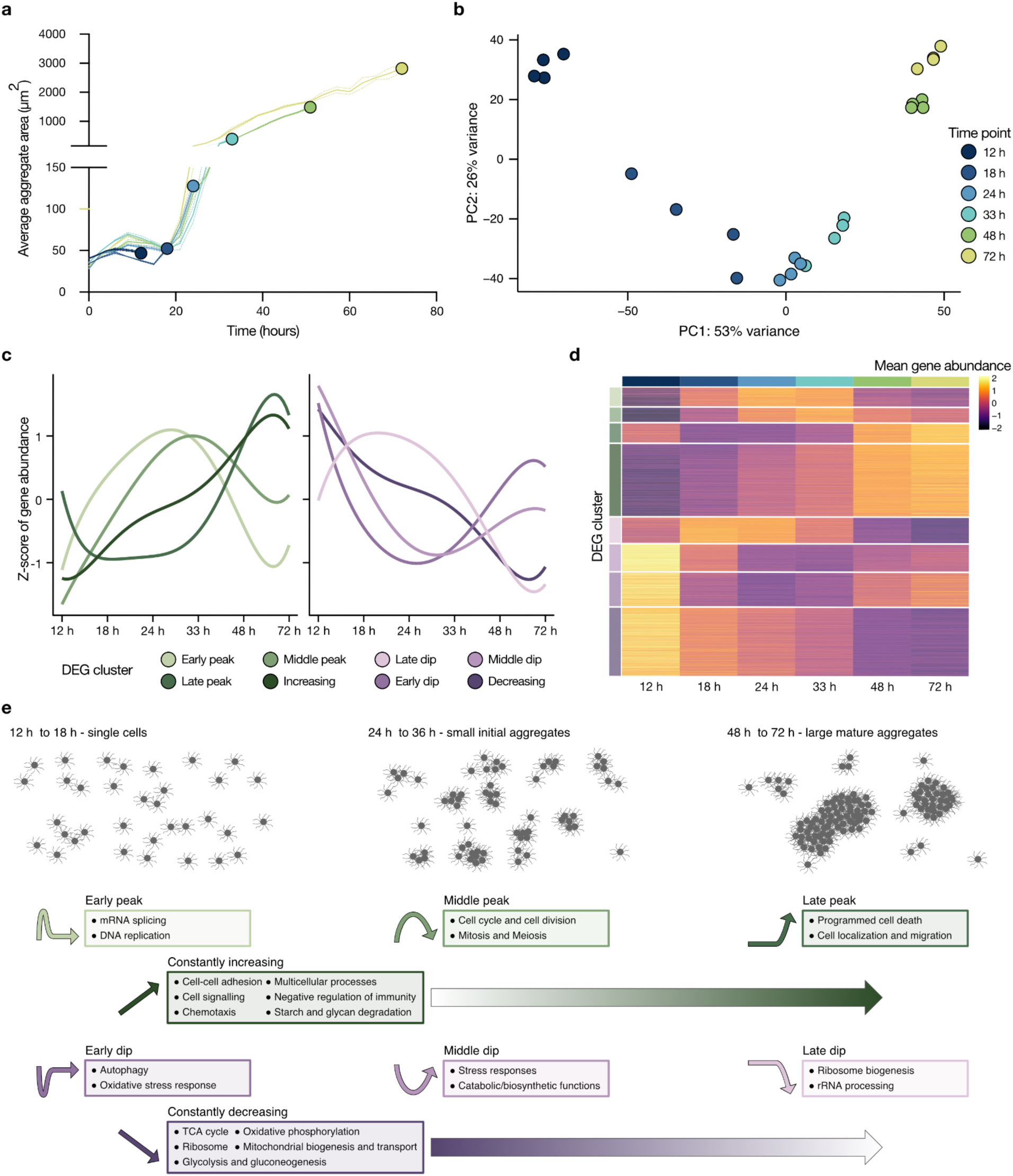
Dynamics of gene expression during *M. vibrans* aggregation time series. a) Average aggregate area in replicate *M. vibrans* monoxenic cultures used for assessing gene expression taken at 12, 18, 24, 33, 48, and 72 h post-inoculation in fresh media. The coloured circle represents the final time point when RNA was harvested for that set of replicates. b) PCA plot of gene expression profiles from replicate cultures taken at the different points in the time series showing the variance between replicates and time points. c) The average Z-score gene expression trends for eight differentially expressed gene (DEG) clusters that arose from expression profile clustering analysis. d) Heatmap of mean gene abundances across the time series for all genes assigned to each DEG cluster. Mean gene abundances were calculated across replicates from each time point and centered and scaled using the variance-stabilizing transformation of normalized gene counts. e) Schematic overview of the main groups of genes that increased and decreased in expression across the time series from each DEG cluster. See also Extended Data Figs. 5-8, Figs. S1-2, and Data S8-9. The constantly increasing and constantly decreasing DEG clusters are also referred to without “constantly”.

To determine which genes drive these shifts during *M. vibrans* aggregation, we identified significantly differentially expressed genes (DEGs) across the time series (see Methods; Data S8). Of these, 4884 genes (40.27 % of *M. vibrans* genes) formed eight DEG clusters with overarching patterns in gene expression (Fig. 4c–d, and Extended Data Fig. 5c, 6a, and 7a, and Data S8). Four of these DEG clusters are composed of net upregulated genes: early peak (n = 326), middle peak (n = 250), late peak (n = 334), and constantly increasing (n = 1268). While another four DEG clusters are composed of net downregulated genes: late dip (n = 445), middle dip (n = 470), early dip (n = 591), and constantly decreasing (n = 1200).

We then examined the gene functions enriched in each DEG cluster using Gene Ontology (GO) terms, Kyoto Encyclopedia of Genes and Genomes (KEGG) pathways, and Pfam protein domain gene annotations (Fig. 4e, Extended Data Fig. 6b, 7b, and 8, Fig. S1-2, Data S9, and Supplementary Discussion 1).^35–37^ Before aggregation, central metabolic, biosynthetic, and stress-response functions became downregulated and continued to decrease in gene expression, likely after an initial period of energy gain and growth. Subsequently, the formation of small aggregates was directly preceded by an initial burst in transcription, translation, protein turnover, mRNA splicing, and DNA replication. Mitosis, cell division, and (unexpectedly) meiosis were upregulated directly after the formation of small aggregates, suggestive of sexual reproduction. Remarkably, multicellularity-related functions (e.g., cell adhesion and signalling) continually increased in gene expression as aggregates first formed and then matured, suggesting a key role in *M. vibrans* aggregation. Simultaneously, functions suggestive of the uptake of bacterial prey (related to phagocytosis and negative regulation of immunity) were upregulated. Finally, after the formation of large mature aggregates, cell localization and programmed cell death-related functions were upregulated, suggesting the potential for cell rearrangements within aggregates.

Together these enrichment analyses suggest that key functions relevant to animal origins are involved in *M. vibrans* aggregation. For example, the early upregulation of mRNA splicing functions indicates alternative splicing, which is thought to have provided a source of functional innovation for evolving multicellularity in animals.^38^ In addition, the enrichment of meiosis-related functions during *M. vibrans* aggregation is indicative of sexual reproduction, whose integration with multicellular life stages is thought to have been an important step in animal origins. Sexual reproduction has previously been shown to occur in the small aggregates (swarms) formed by *S. rosetta* in response to the bacterium *Vibrio fischeri* and in response to nutrient limitation.^3,39^ Furthermore, cell adhesion, signalling, and transcriptional regulation are critical in animal multicellular development.^40^ Thus, we next sought to investigate the repertoire of adhesion, signalling, and transcriptional regulation genes important for animal development found in *M. vibrans* and further assess their individual expression profiles.

### Genes key to animal multicellularity are expressed during *M. vibrans* aggregation

We identified 489 genes in *M. vibrans* involved in multicellularity and development in animals, and that could be confidently linked to cell adhesion (n=73), cell signalling (n=363), and transcriptional regulation (n=53) (Data S10). Of these, 307 changed significantly in expression across the *M. vibrans* aggregation time series, with 89%, 76%, and 55% of cell adhesion, cell signalling, and transcriptional regulation related genes, respectively, assigned to net upregulated DEG clusters (Fig. 5a). Many of these genes are critical for animal multicellularity. Some were previously thought to be animal-specific, but are now known to have deeper evolutionary roots, including in the uropisthokont, urholozoan, and urfilozoan ancestors (Fig. 5b–c, and Data S10).^1,12,41–43^

**Figure 5.**
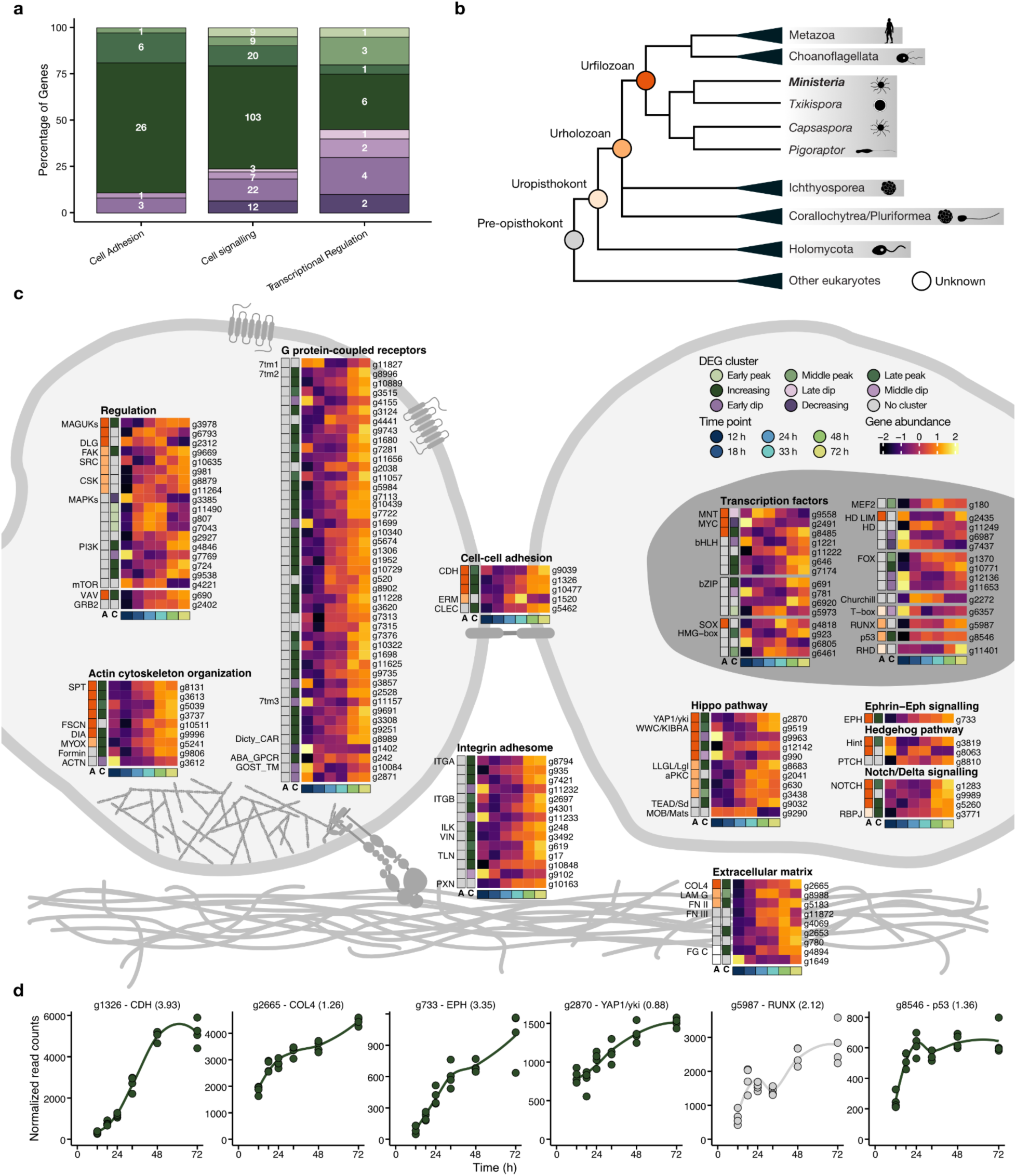
Key animal multicellularity genes are expressed during *M. vibrans* aggregation. a) Percentage of genes linked to cell adhesion, cell signalling, and transcriptional regulation assigned to each differentially expressed gene (DEG) cluster. b) Overview of Opisthokonta species relationships and ancestors indicated by coloured circles. c) Schematic of two *M. vibrans* cells and heatmaps showing the expression patterns of different categories of multicellularity-related genes that are significantly differentially expressed across the aggregation time series. The bottom row indicates the time point according to the legend, the first column (column “A”) indicates the predicted gene age according to the coloured ancestors in panel b, and the second column (column “C”) indicates DEG cluster assignment. Mean gene abundances were calculated across time point replicates and centered and scaled using the variance-stabilizing transformation of normalized gene counts. Spectrin; SPT, fascin; FSCN, diaphanous; DIA, myosin X; MYOX, formin-like protein; Formin, alpha actinin; ACTN, alpha type IV collagen; COL4, laminin G; LAM G, fibronectin type II and type III; FN II and III, fibrinogen C-terminal; FG C, integrin alpha; ITGA, integrin beta; ITGB, integrin-linked kinase; ILK, alpha-catenin/vinculin family; VIN, talin; TLN, paxillin; PXN, cadherin; CDH, ezrin-radixin-moesin family; ERM, C-type lectin; CLEC, membrane-associated guanylate kinases; MAGUKs, discs large MAGUK; DLG, focal adhesion kinase; FAK, tyrosine-protein kinase c-Src; SRC, C-terminal Src kinase; CSK, mitogen-activated protein kinases; MAPKs, phosphatidylinositol 3 kinase; PI3K, mechanistic target of rapamycin; mTOR, Vav guanine nucleotide exchange factor; VAV, growth factor receptor-bound protein 2; GRB2, rhodopsin/class A; 7tm1, adhesion/secretin/class B; 7tm2, glutamate/class C; 7tm3, cAMP/classE; Dicty_CAR, abscicic acid receptor; ABA_GPCR, GOST seven transmembrane; GOST_TM, yes-associated protein 1/yorkie; YAP1/yki, WW and C2 domain containing/kidney and brain expressed protein; WWC/KIBRA, lethal 2 giant larvae; LLGL/Lgl, protein kinase C; aPKC, TEA domain transcription factor/scalloped; TEAD/Sd, MOB kinase activator/mob as tumor suppressor; MOB/Mats, eph receptor; EPH, Hint/Hog domain found in Hedgehog; Hint, patched; PTCH, notch; NOTCH, recombining binding protein suppressor of hairless; RBPJ, basic helix loop helix; bHLH, MAX network transcriptional repressor bHLH; MNT, MYC proto-oncogene bHLH; MYC, basic leucine zipper; bZIP, high mobility group box; HMG-box, SRY-related HMG-box; SOX, myocyte-specific enhancer factor 2 MADS-box; MEF2, homeodomain; HD, forkhead; FOX, runt-related; RUNX, rel homology DNA-binding; RHD. d) Plots of normalized gene counts across the aggregation time series for selected upregulated genes with diverse functions and filozoan or holozoan origins. Log2 fold change between 12 and 72 h is indicated in brackets. See also Extended Data Fig. 9, Fig. S3, and Data S10.

First, we observed significant upregulation of several adhesion-related genes, including homologs responsible for building cellular scaffolding (i.e., actin cytoskeleton organization), the extracellular matrix (ECM), cell-ECM adhesion, and cell-cell adhesion in animals (Fig. 5, Extended Data Fig. 9, Fig. S3, Data S10, and Supplementary Discussion 2). These genes include spectrin, fascin, and diaphanous gained in the urfilozoan ancestor, and myosin X gained in the urholozoan ancestor, which in animals function in actin filaments, epithelia, and filopodia formation.^11,44,45^ Diverse ECM-related genes were also upregulated, including alpha type IV collagen–thus far the only case outside Metazoa.^38^ The integrin adhesome has pre-Opisthokonta origins and is central in animal cell-ECM adhesion.^46–49^ Most components were upregulated together with holozoan-specific kinases that function in integrin-mediated signalling. Upregulated genes involved in animal cell-cell adhesion included a putative C-type lectin (not previously found in Filasterea) and a homolog of the Holozoa-specific ezrin-radixin-moesin family.^17,50,51^ In addition, we found that *M. vibrans* encodes seven cadherin homologs, several of which were upregulated and increased up to 15 times in gene expression. Cadherins are Filozoa-specific with only one in *C. owczarzaki*.^47,52^

We next observed the activation of numerous cell signalling pathways critical in animals (Fig. 5, Extended Data Fig. 9, Fig. S3, Data S10, and Supplementary Discussion 2). These signalling mechanisms coordinate animal cell behavior and ensure the formation of a cohesive unit, indicating a potential for coordination and communication between aggregating *M. vibrans* cells. Many tyrosine kinases (TKs), mitogen-activated protein kinases, and phosphatidylinositol-3-kinases were differentially expressed, alongside membrane-associated guanylate kinases, which are Filozoa-specific and regulate animal cell-cell adhesion.^53,54^ Small GTPases had variable expression patterns: the majority of guanine nucleotide exchange factors, including the filozoan-specific VAV,^55^ GTPase-activating proteins, and the growth factor receptor-bound protein 2 signalling adaptor were upregulated, while guanine nucleotide dissociation inhibitors were downregulated. In addition, the majority of differentially expressed G-protein coupled receptors (GPCRs) were upregulated. These particularly included adhesion/secretin/class B GPCRs, the number of which encoded by *M. vibrans* is exceptional outside of Metazoa.^56^ A putative Filozoa-specific eph receptor^57^ from the Ephrin-EPH pathway critical for animal morphogenesis) increased over 10 times in gene expression. All potential Notch receptor homologs, which again are Filozoa-specific,^29,43,58^ were also upregulated. Furthermore upregulated was a transcription factor (TF) involved in Notch/Delta signalling: recombining binding protein suppressor of hairless. The Hippo pathway regulates animal tissue size. Most of its identified components were differentially expressed, including the Filozoa-specific Yorkie (Yes-associated protein 1).^59^ The Hedgehog pathway regulates cell differentiation, and most of its components are animal-specific.^60^ However, we identified several conserved genes encoding hedgehog (hh) hint/hog domains that were differentially expressed,^29,52,61^ along with the hedgehog receptor patched.

Finally, we found that a large collection of TFs is differentially expressed in *M. vibrans*, including many involved in transcriptional regulation during animal development (Fig. 5, Extended Data Fig. 9, Fig. S3, Data S10, and Supplementary Discussion 2). In contrast to cell adhesion and signalling, a nearly equal number of differentially expressed transcription factors were net up and downregulated across the *M. vibrans* aggregation time series. The individual expression profiles of these TFs were diverse, revealing a complex regulatory system with which to deploy multicellular genes in *M. vibrans*. A number of differentially expressed *M. vibrans* TFs are relatively recent innovations. T-box and Rel homology DNA-binding TFs were gained in the uropisthokont ancestor, runt-related and p53 TFs in the urholozoan ancestor, and Myc-Max network, homeodomain LIM, and SRY-related HMG-box (SOX) TFs in the urfilozoan ancestor.^46,62–65^ The expression of these TFs during *M. vibrans* aggregation suggests that they were already regulating multicellularity in unicellular animal ancestors.

We note that some sets of animal multicellularity-related genes have previously been found upregulated in different cell types of diverse unicellular animal relatives.^11,66,67^ Yet, *M. vibrans* is exceptional in expressing homologs of nearly the full complement of such genes during aggregation. Although we do not know the function of most of these genes in a non-animal context, several have been found to have analogous functions relevant for animal multicellularity.^11,65,68,69^

Altogether, we found that components of most cell adhesion and cell signalling pathways found in *M. vibrans* were differentially expressed and upregulated across the aggregation process. Homologs of the various TFs involved in animal development were also differentially expressed with diverse expression patterns indicating coordination of various phases of the *M. vibrans* aggregation process. Moreover, gene homologs with recent evolutionary origins in Opisthokonta, Holozoa, and Filozoa were differentially expressed and most often upregulated. Our results show that an extensive gene repertoire key in animals is also involved in the development of *M. vibrans* aggregates. Thereby suggesting conservation of mechanisms for cell adhesion, signalling, and transcriptional regulation between *M. vibrans* and animal multicellularity.

### Improved feeding and meiosis are potential drivers of *M. vibrans* aggregation

Finally, we asked what advantage aggregation affords *M. vibrans*. For unicellular organisms, aggregation can have multiple benefits, including improved feeding or mating.^70–72^ *M. vibrans* is a known bacteriovore,^28^ and our monoxenic culture exhibited clear phagocytosis of the *T. lucentensis* food bacteria (Fig. 6a–c). Furthermore, we observed that the aggregating monoxenic culture grew better than the non-aggregating xenic culture (Fig. 2a). The correlation between growth and aggregation led us to hypothesize that aggregation promotes *M. vibrans* replication. To test this hypothesis, we prevented aggregate maintenance through physical disruption (Extended Data Fig. 10a). We then asked if disaggregated cells grew worse than those that remained in aggregates. Indeed, disaggregated cells replicated worse than those left undisturbed (Fig. 6d). Therefore, our findings are consistent with aggregation promoting *M. vibrans* growth, although we cannot rule out other possible effects of the physical disruption of aggregates.

**Figure 6.**
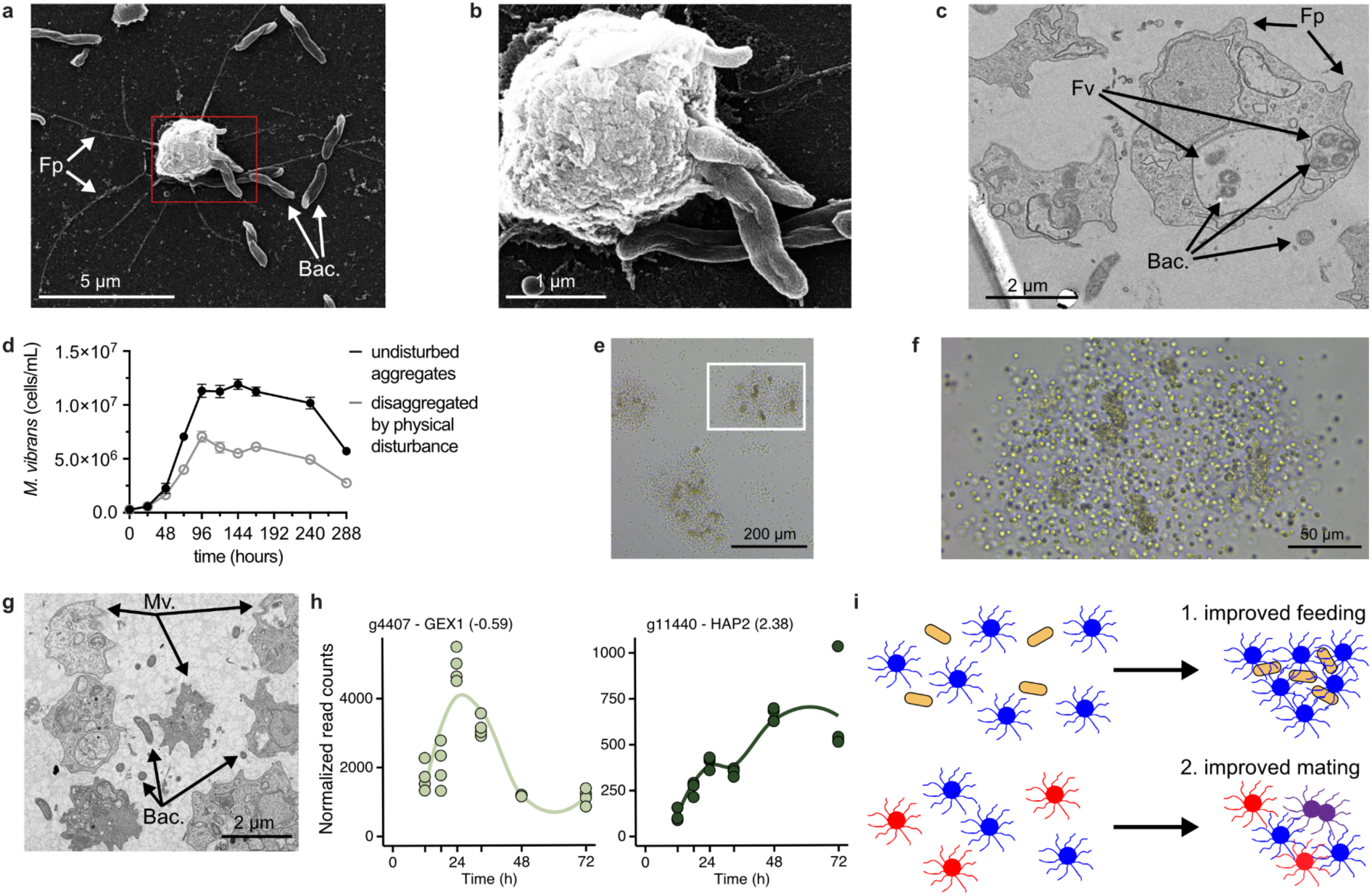
Evolutionary drivers of aggregation. a) Scanning electron micrograph revealing *M. vibrans* engulfing *T. lucentensis* bacteria (Fp = filopodia, Bac = bacteria). b) Expansion of the red box in panel (a). c) Transmission electron micrograph revealing *T. lucentensis* bacteria within *M. vibrans* vacuoles (Fp = filopodia, Bac = bacteria, Fv = food vacuole). d) Cell counts of *M. vibrans* in cultures with/without physical disruption of aggregates. Error bars represent SE of a biological triplicate. e) Brightfield microscopy image of *M. vibrans* aggregates with putative bacterial biofilm material embedded within. f) Expansion of the white box in panel (e). g) Transmission electron micrograph revealing *T. lucentensis* bacteria in the intercellular space of an *M. vibrans* aggregate (Mv = *M. vibrans* cell, Bac = bacteria). h) Normalized gene expression counts of GEX1 and HAP2 gene homologs, which are meiosis-specific marker genes, during the time course of aggregation. Log2 fold change between 12 and 72 h is indicated in brackets. i) Scheme illustrating potential benefits of aggregative multicellularity for *M. vibrans* (and its ancestors): improved feeding and mating. See also Extended Data Fig. 10 and Fig. S5.

A possible explanation for the improved replication is an increase in food uptake. Cells in the middle of an aggregate may not have access to bacteria, which would actually inhibit feeding. However, as aggregates form and morph, they may also entangle bacteria within their community, providing more efficient feeding. In support of this hypothesis, we have observed material embedded within aggregates that resembles bacterial biofilms (Fig. 6e–f). Electron microscopy further revealed bacterial cells within the intercellular space of *M. vibrans* aggregates (Fig. 6g). We have also observed that the number of *T. lucentensis* food bacteria in the culture and number of mapped reads rose with the formation of small aggregates before dropping shortly after *M. vibrans* formed large aggregates (Extended Data Fig. 10b and Fig. S4). The formation of aggregates and drop in *T. lucentensis* count also correlated with the expression of phagocytosis-associated genes in *M. vibrans*, with initially high expression of the Arp2/3 complex, followed by later upregulation of actin cross-linking and remodelling proteins (Extended Data Fig. 10c–d, Fig. S5, and Supplementary Discussion 3). Further work is required to fully test this hypothesis, but aggregation may improve the efficiency of bacterial capture by *M. vibrans*.

Additionally, aggregates may promote opportunities for mating between cells, as shown in diverse organisms, including the choanoflagellate *S. rosetta*.^3,39,73,74^ Mating can improve fitness by transmitting beneficial genetic traits, spawning beneficial combinations of genes,^75^ and providing adaptive flexibility.^76^ Indeed, we found that multiple genes associated with meiosis, including meiosis-specific markers GEX1 and HAP2,^77–79^ were upregulated during aggregation, particularly in small aggregates (Fig. 6h, Extended Data Fig. 10d, Fig. S5, and Supplementary Discussion 3), suggesting another plausible benefit for aggregation. Overall, our results indicate improved feeding and increased mating within *M. vibrans* aggregates. Therefore, these increased behaviors also may have been evolutionary drivers benefitting aggregative multicellularity in the unicellular ancestors of animals (Fig. 6i).

## Discussion

In our study we have demonstrated aggregative multicellularity in the filasterean *M. vibrans*. Taken together with observed cellular aggregation in the distant filasterean *C. owczarzaki*,^11^ two choanoflagellates,^3,4^ and multiple extant animals,^80–83^ we can thus infer that aggregation was ancestral and potentially employed by the unicellular ancestors of animals. Furthermore, we found that *M. vibrans* expresses many animal multicellularity genes while it forms large stable aggregates. A new analysis of chemically regulated aggregation in *C. owczarzaki* reveals a parallel correlation between aggregation and dynamic multicellular gene expression.^84^ In both cases, Holozoa-specific genes that drive cell adhesion, signalling, and gene expression in animals are upregulated in aggregates. Since such conserved genes are involved in holozoan cellular aggregation, our results suggest that the common ancestor of filastereans and animals had the capacity to use these genes for aggregative multicellularity.

Apart from the filastereans *M. vibrans* and *C. owczarzaki*, many other protists exhibit aggregative behaviors (e.g., *Fonticula*, *Acrasis*, *Dictyostelium*, choanoflagellates, Rhizaria 2012).^3,4,85–89^ In fact, we suspect this behavior is even more common across protists, but the environmental triggers have yet to be reported (as was the case with *M. vibrans* and *C. owczarzaki* until we discovered the right inducing conditions). The widespread nature of aggregative behavior suggests major fitness benefits and supports the hypothesis that the ancestor of animals was capable of aggregation. Several potential benefits have been reported for simple multicellularity.^5,90,91^ Our initial observations with *M. vibrans* suggest that aggregation may improve mating or feeding behaviors. We observed several meiosis genes upregulated during aggregation. Therefore, like many organisms (including choanoflagellates)^3^ *M. vibrans* may leverage aggregation to improve mating efficiency. Alternatively, aggregation improved *M. vibrans* replication, and bacteria were trapped within the aggregates. Entrapment of bacteria may increase *M. vibrans* fitness. Initially, the aggregation of cells through a web of interconnected filopodia could improve the capture of bacteria. Subsequently, storing bacteria within the intercellular aggregate space may provide sustained food as the environment changes and bacteria become less abundant. Perhaps bacteria may even replicate within the aggregates, providing a constant source of food. The long-term stability of the aggregates is consistent with this hypothesis. This behavior of harboring bacteria resembles the primitive ‘farming’ behavior in *Dictyostelium*, which collects bacteria within its multicellular fruiting bodies.^92^ Some choanoflagellates also are known to harbor bacteria within their multicellular clusters.^93^ The green alga *Volvox carteri* and the ciliate *Stentor coeruleus* also form multicellular bodies to enhance feeding, thus accelerating replication.^94–96^ Of course, many extant animals have also evolved diverse mechanisms for maintaining microbiomes, and microbial cues and interactions play important roles in animal development.

Despite the prevalence of multicellular aggregation across numerous protists and diverse animal cell types, aggregation is rarely considered to be a key trait in the first multicellular animals. A primary concern is the potential for genetic conflict within aggregates of non-clonal cells.^90,97^ However, recent work has refuted the concerns over cheating: spatial structure combined with group selection facilitates the survival of cooperators against cheaters in an aggregate.^23^ Moreover, our study demonstrates that *M. vibrans* aggregates are stable once formed, persisting over two months. In other well-studied cases of aggregative multicellularity such as sorocarps, aggregation is a short-lived temporary life stage in response to environmental stress prior to fruiting-body formation and spore release.^88,98–100^ Stable aggregation would have allowed more opportunities for complex multicellularity to evolve.

Another concern with the importance of aggregation for the origin of animals was a lack of clear homology between aggregation-inducing genes in protists and in animals. The best-studied aggregating protists are only distantly related to animals.^88,100–105^ Therefore, their aggregation behaviors lack involvement of Holozoa-specific genes that are involved in animal multicellular behaviors. Now this work reveals conservation between genes involved in multicellularity of animals and cellular aggregation of their closest relatives. This finding suggests that the ancestor of animals plausibly used these genes for reversible multicellular aggregation. In other words, the origin of those genes that later on were crucial for animals to evolve could be related to cell aggregation. In fact, the most morphologically similar aggregative behavior in an extant animal is exhibited by sponge cells, which also upregulate genes involved in cell adhesion, integrin, and actin remodeling.^106,107^ Furthermore, wound healing responses (which involve the reaggregation of cells and maturation into a tissue) in diverse animals (e.g., sponges, cnidarians, and ctenophores) share transcriptional similarities with *M. vibrans* aggregation.^80–83^ The model sponge *Sycon ciliatum* upregulated genes involved in actin filaments and signal transduction during wound healing.^80^ The model cnidarian *Nematostella vectensis* also expressed similar adhesion and signalling genes related to cell-substrate adhesion, integrin, GPCRs, and receptor tyrosine kinases.^81,82^ Similarly, the ctenophore *Mnemiopsis leidyi* expressed cytoskeletal remodeling and signalling genes related to GPCRs and GTPases.^83^ These similar expression profiles of adhesion, cytoskeleton, and signalling genes in *M. vibrans* aggregation and animal wound healing support the hypothesis that simple cellular aggregation was a precursor to more complicated cellular reorganization processes in the first animal. This hypothesis does not necessarily exclude the involvement of the other two main types of multicellular behavior shared between unicellular holozoans and animals (incomplete separation after clonal division and coenocytic development). All three behaviors may have been important in the life cycle of the ancestor of animals, since each behavior presents a unique advantage to an organism with a complex lifestyle (clonal division avoids cheating, aggregation provides rapid changes and dynamic structures, and coenocytic division yields different cell types).^1^

In summary, we have here uncovered how a close unicellular relative of animals robustly forms homogeneously large multicellular aggregates. It employs many genes that animals use for cell adhesion, cell signalling, and transcriptional regulation through the process of aggregation. Therefore, we hypothesize that the filozoan ancestor of animals used newly acquired ‘multicellularity’ genes for multicellular aggregation behavior, perhaps gaining an edge in acquiring food or weathering environmental challenges. The origin of those multicellular genes together with the capacity to form stable aggregates using those genes, may have pre-adapted those ancestors to eventually evolve into the first multicellular animals.

## Methods

### Culturing *M. vibrans*

*M. vibrans* ATCC 50519 (primary strain in this study) and *M. vibrans* DSM 114853 were cultivated with 5% seawater complete medium (SWC, 32.9 g/l tropic marin salt, 5 g/L peptone, 3 g/L yeast extract and 3 ml/L glycerol, pH 8.0) in vented T-25 flasks (Fisher Scientific, Cat #: FB012935) at room temperature (23 °C). They were subcultured every 3 days into a fresh T-25 flask by splitting cultures 1:10 with fresh 5% SWC medium.

### 16S rDNA metagenomic sequencing

Metagenomic DNA of *M. vibrans* ATCC 50519 xenic culture was extracted with Invitrogen™ PureLink™ Microbiome DNA Purification Kit (ThermoFisher, Cat. # A29790). Metagenomic 16S rDNA sequencing was performed by Molecular Research LP (USA) using bTEFAP® illumina MiSeq with primer 515F/806R. Comprehensive taxonomic analysis zOTUs percentage and counts compilations at each taxonomic level were performed.

### Monoxenic *M.vibrans* cultures

*M. vibrans* Tong ATCC 50519 xenic culture was treated with antibiotic cocktail (ofloxacin 10 µg/ml, kanamycin 50 µg/ml, streptomycin 50 µg/ml, ampicillin 50 µg/ml and chloramphenicol 68 µg/ml) in artificial seawater (ASW) to kill environmental bacteria. After 48 hours, cells were pelleted and resuspended with fresh ASW containing antibiotics for another 48 hours. 100 *M. vibrans* cells were then sorted by flow cytometry LSRII into wells of a 24-well plate containing 500 µl ASW with antibiotics to avoid bacterial contamination. Cells were maintained in the dark at RT overnight. After checking the wells with a microscope (Leica DMIL LED) the wells without visual bacteria were supplemented with 2 ml ASW and 10 µl of *Thalassospira lucentensis* bacteria. Recovered *M. vibrans* cells were transferred to T-25 vented flasks containing 5% SWC after one week. Metagenomic DNA of putative monoxenic *M. vibrans* culture was extracted with Invitrogen™ PureLink™ Microbiome DNA Purification Kit (ThermoFisher, Cat. # A29790). Metagenomic 16S rDNA V4 amplicon was sequenced with primer 16S V4 forward and reverse (515F & 806R) to confirm the monoxenic culture of *M. vibrans* in Center for Genomics and Bioinformatics, Indiana University Bloomington (IUB-CGB). Libraries were prepared with NEXTFLEX16S V4 Amplicon-Seq Kit 2.0 and DNA was purified with Agencourt AMPure XP Magnetic Beads. Paired-end sequencing was performed on a MiSeq Nano v2 (500 cycles). The data was demultiplexed using bcl2fastq, version 2.20.0, and analyzed by Marker Based Population Analysis (16S). The *M. vibrans* monoxenic culture derived from DSM 114853 was generated with the same above protocol. The culture was then confirmed as monoxenic by metagenomic 16S rDNA sequencing in Molecular Research LP (USA) with the same procedure applied for the *M. vibrans* ATCC 50519 xenic culture. SRA accessions for all 16S rDNA amplicon sequence reads can be found in Data S5.

### Growth curve of each *M. vibrans* cell line

Each *M. vibrans* cell line was cultivated with their own 5% seawater complete medium (SWC) in a 24-well plate at RT. *M. vibrans* cells were counted with a hemocytometer every 24 hours after pipetting to disperse the possible cell clumps.

### Quantification of aggregation area of each *M. vibrans* cell line

200 μl of each *M. vibrans* cell line (5.0 x 10^5^ cells/ml) was placed into ultra-low attachment 96-well plate (VWR, Cat. # 29443-034) in fresh 5% SWC for imaging with Cytation C10 imager (Agilent BioTek) at 4x objective every 3 hours.

### Scanning electron microscopy (SEM)

500 µl *M. vibrans* cells were placed into poly-L-lysine coated round coverslips (VWR, Cat # BD354085), and the cells settled for 1 hour. 200 µl media was removed, and the cells were washed gently with 200 µl artificial sea water (ASW) three times. Two fixation steps were applied to the cells. In the first step, the cells were fixed for 30 minutes 200 µl 2% glutaraldehyde in 0.1M sodium cacodylate buffer, pH 7.4 (Electron Microscopy Sciences, Cat #:16536-15), after first washing them twice in this solution. In the second step, cells were fixed for 30 minutes with 1% osmium tetroxide (Sigma, Cat # 75632-10ML). Continuous dehydration was performed by washing the cells twice with an ethanol gradient (30%, 50%, 70%, 80%, 90%, 95% and 100%), soaking the cells in each solution for 10 min before moving to the next. Samples were critical-point-dried in liquid CO_2_ using a samdri-790 critical-point drying apparatus. They were subsequently put on SEM aluminum mount for amray 1400 (Electron Microscopy Sciences, Cat. # 75174) with carbon adhesive tabs (Electron Microscopy Sciences, Cat. # 77825-12), sputter-coated with gold-palladium (80:20) with safematic CCU-010. Samples were imaged with an FEI Quanta 600 Scanning Electron Microscope (Indiana University Bloomington).

### Transmission electron microscopy (TEM)

*M. vibrans* cell aggregates in ultra-low attachment 96-well plates were washed with ASW to remove extraneous material. Wells were filled with 2.5% glutaraldehyde and 2.0% osmium tetroxide in 0.1 M sodium cacodylate buffer, pH 7.2 (>200 µl) on ice after removing ASW. Cells were washed with 0.1 M sodium cacodylate buffer (pH 7.2) four times and then were embedded with 1% agarose in 0.1 M sodium cacodylate buffer. Cells in agarose plugs were dehydrated in a graded ethanol series of 30%, 50%, 75% with 2% uranyl acetate, 90%, 95% and 100% for 10 minutes each. The ethanol solutions were kept on ice. Samples were placed in a 1:1 mixture of ethanol and acetonitrile for one hour and then in 100% acetonitrile for another hour. The samples were placed in a mixture of acetonitrile and Embed 812 (Electron Microscopy Sciences, catalog number 14130) at the ratios of 3:1, 1:1, and 1:3 for two hours at each dilution, and then in 100% Embed 812 three times, one time overnight and the other two times for a minimum of two hours. The resin-embedded sample was polymerized at 65 °C for 18 hours. The resin blocks were sectioned at 70 nm using a Leica Ultracut UCT ultramicrotome. Ultrathin sections, stained with uranyl acetate and lead citrate, were imaged with a JEOL JEM 1010 transmission electron microscope at 80 kV using a Gatan 794 camera.

### Blocking replication of *M. vibrans*

2 ml *M. vibrans* cells (1.0 x 10^5^ cells/ml) in fresh 5% SWC in a 24-well plate were treated with 20 μg/ml aphidicolin (Sigma, Cat. # 178273-1MG) or DMSO. Cell density was counted every 24 hours with a hemocytometer for 6 days to determine the concentration of aphidicolin that can block cell replication. Once these conditions were established, 200 µl of 20 μg/ml aphidicolin- or DMSO-treated *M. vibrans* cells (5.0 x 10^5^ cells/ml) were placed into an ultra-low attachment 96-well plate for imaging with a Cytation C10 imager every 2 hours for 3 days.

### *T. lucentensis* whole genome sequencing

*T. lucentensis* was cultivated in SWC medium at 30 °C at 220 rpm overnight. 1 ml *T. lucentensis* cells were used for genomic DNA extraction with the Wizard genomic DNA isolation kit (Promega, Cat. # PAA1120). The genomic DNA of *T. lucentensis* was sent to Eurofins for whole genome sequencing. Rapid Barcoding Kit V14 was used for the library preparation. The library was then sequenced with Oxford Nanopore GridION by using Kit 14 and the R10.4.1 nanopore. The genome was annotated with Bakta v1.9.2. The SRA accession can be found in Data S5.

### RNA-Sequencing

Total RNA extraction was modified from Ocaña-Pallarès^108^ Briefly, 1.0 x10^7^ cells were collected into 15 ml falcon centrifuge tube after incubating in ultra-low attachment plates in a Cytation C10 imager at 26 °C for 12, 18, 24, 33, 48 and 72 hours (4 replicates each). Supernatants were removed by pelleting at 4000 xg for 5 minutes. Cells were lysed with 1 ml Trizol for 5 minutes and 200 µl 1-bromo-3-cloropropane for 15 minutes at RT. Total RNA and DNA then were precipitated with isopropanol and rinsed with fresh-prepared 75% ethanol. The pellet was redissolved in 50 µl of 1x Dnase I buffer, and DNA then was digested by DNase I (ThermoFisher, Cat. # EN0521,) and RNAseOUT (ThermoFisher, Cat. # 10777019). The RNA was purified by precipitation with 5 µl LiCl [5M] in x2 total volume (110 µl) 100% ethanol (previously cooled at −20°C). RNA was washed with 70% ethanol (previously cooled at −20°C) and dissolved in 20 µl nuclease free water. Samples were sent to NOVOGENE for mRNA library preparation (poly A enrichment) by using poly-T oligo-attached magnetic beads. Quantified libraries were sequenced on Illumina platform NovaSeq X Plus Series PE150. SRA accessions for RNA-sequence reads can be found in Data S5.

### *M. vibrans* functional annotation

All analyses were performed using the recently published *M. vibrans* ATCC 50519 genome.^34^ Genomic data files available from the figshare^109^ repository associated with the publication were used for all analyses, specifically the files “Mvib.PASAupdated_BRAKERproteins.070617.fa”, “Mvib.PASAupdated_BRAKERproteins.070617.gff” for protein sequences and gene annotations, and the file “Mvib_cleangenome.withMito18S.plusaddedfromdraft.300417.fa” for nucleotide sequences. Genes were functionally annotated using several different tools and databases. Emapper v.2.1.9^110^ with eggNOG v.5.0.0^111^ with the settings “-m hmmer --tax_scope auto --tax_scope_mode inner_narrowest --go_evidence all --pfam_realign denovo” at both the “Eukaryota” and “Metazoa” levels. InterProScan v.5.62-94.0^112^ with the settings “-appl Pfam, TIGRFAM, PANTHER, -iprlookup, -pa”. Top hits using BLAST+ v.2.14.1+^113^ blastp with the settings “-max_target_seqs 5, -max_hsps 1” against the *C. owczarzaki* genome (GCF_000151315.2), *Homo sapiens* GRCh38 genome (GCF_000001405.40), and EukProt V3 TCS dataset.^84^ DIAMOND v.2.1.9^114^ with the options “--max-target-seqs 5, --more-sensitive” against Swiss-Prot,^115^ and NCBI’s RefSeq and NR databases.^116^ BlastKOALA v.3.0^117^ with the KEGG GENES database “Eukaryotes”. Functional annotations were combined into an overview across genes using a custom script “make_Mvib_annotations_table.py” (Data S6).^118^

### Processing of RNA-sequencing reads

RNA-sequencing reads were processed and mapped against the *M. vibrans* genome using the nf-core/rnaseq pipeline v.3.14.0^119–121^ with the options “--igenomes_ignore --genome null --skip_bbsplit false - -remove_ribo_rna --aligner star_rsem --stringtie_ignore_gtf --bam_csi_index -profile singularity”. More specifically, adaptors were removed and reads trimmed using Trim Galore! with read quality control checked using FastQC. Reads that mapped against the *Homo sapiens* GRCh38 genome and *T. lucentensis* Mv1 genome were removed using BBSplit from BBMap and ribosomal RNA removed using SortMeRNA.^122^ Reads were then aligned to the *M. vibrans* genome using STAR^123^ and quantified using RSEM.^124^ The resulting gene quantification files were used for downstream analyses^118^ and read counts are available in Data S7.

### Sample relationships analyses

Gene quantification files were imported into RStudio v.2023.12.1+402 with R v.4.3.3 using tximport v.1.30.0^125^ and loaded into a DESeq dataset using DESeq2 v.1.42.1.^126^ Genes with low counts, fewer than ten counts in less than four samples, were removed and counts from the remaining genes (10852 of 12127 genes) normalized using “estimateSizeFactors” to allow for between-sample comparisons (Data S7). The resulting normalized counts were then subjected to a variance stabilizing (vsd) transformation to remove dependence of the variance on the mean using vsn v.3.70.0 (Data S7).^127^ A sample distance matrix of the vsd transformed data was then calculated using “dist” and sample relationships visualized through a principal component analysis (PCA) plot using “plotPCA” and a heatmap of sample to sample distances using pheatmap v.1.0.12.^128^

### Identification and clustering of differentially expressed genes

DEGs that differed significantly in gene expression across the time series were then identified using the likelihood ratio test (LRT) with the options “alpha = 0.0000000001 and filterFun=ihw” using IHW 1.30.0^129^ and DESeq2 v.1.42.1.^126^ Genes with adjusted p-value < 1 × 10-10 were retained for downstream analyses (6472 genes; Data S8). DEG clusters with a minimum size of 250 genes were then identified using “degPatterns” from DEGreport v.1.38.5.^130^ This resulted in eight overarching DEG clusters that captured the most common gene expression profiles across the time series (Data S8), and which were visualized with pheatmap v.1.0.12.^128^

### Functional enrichment analysis

Significantly enriched GO terms and KEGG pathways in each DEG cluster were then identified using “enricher” from clusterProfiler v.4.10.1^131^ with default settings (p-value ≤ 0.05) (Extended Data Fig. 8). An organism database including GO terms annotated from across various tools was compiled using AnnotationForge v.1.44.0.^132^ For GO term enrichment analysis, a topGO data object for “Biological Processes” GO terms was then compiled and significance tested using “runTest” with the options “algorithm=’weight01’, statistic=’fisher’” from TopGO v.2.54.0.^133^ Significantly enriched GO terms (weighted Fisher ≤ 0.001; Data S9) were retained and “calculateSimMatrix” from GOSemSim v.2.28.1^134^ used to calculate semantic score similarity between terms with the settings “ont=’BP’, method=’Re’, and keytype = ‘GID’”. Terms were then reduced using “reduceSimMatrix” with a threshold of “0.9”. Resulting GO terms and similarity data were then plotted using Rrvgo v.1.14.2.^135^ R-analyses of RNA-sequencing data are available in an R markdown file.^118^

### Annotation and analyses of genes of interest

Genes involved in animal multicellularity^12,41,42^ and previously investigated in unicellular holozoans were identified in M. vibrans using compiled functional annotation data, the presence of Interpro,^136^ Pfam,^137^ and PANTHER^138^ protein domains, and reciprocal best hits against human homologs using BLAST+ v.2.14.1+ (Data S10).^113^ Previously investigated phagocytosis-related genes^11,139^ and meiosis-related genes^140,141^ were identified on the basis of the presence of conserved protein domains and functional annotation data (Data S11). Individual gene expression profiles were visualized in heatmaps using pheatmap v.1.0.12.^128^

### Gene age inference

Gene ages for key multicellularity genes were inferred using several different strategies. First, gene family age based on originations from a recent ancestral state reconstruction of Opisthokonta genome evolution were collected (REF).^34^ Second, gene age was inferred using GeneBridge v.0.99.2.^142^ Here, the species topology and gene families from Ocaña-Pallarès et al. were used^34^ and the tree uploaded using ape v.5.8.1 (REF).^143^ GeneBridge was run with a threshold of “0.3” and “100000” permutations. Third, results from prior studies investigating the evolutionary history and distribution of these genes were compiled.^38,43,44,46,50–53,55–58,60,62,65,144–152^ A consensus gene age was then selected, with inferences from prior studies taking precedence (Data S10).

## Data visualization

In addition, for various R analyses, data was formatted using tidyverse v.2.0.0,^153^ dplyr v.1.1.4,^154^ and plyr v.1.8.9,^155^ with additional plots visualized using ggplot2 v.3.5.2^156^ and cowplot v1.1.3,^157^ together with colour palettes from RColorBrewer^158^ and viridis v.0.6.5.^159^

## Data availability

Sequencing reads have been deposited at the NCBI SRA database under BioProject PRJNA1229532 for RNA-sequencing (SRR32514592 to SRR32514615), 16S rDNA amplicon sequencing (SRR33469057 to SRR33469059), and the *Thalassospira lucentensi*s strain Mv1 (SRR33469287). Information about which SRA accession corresponds to which sample alongside BioSample accessions can be found in Data S5. The assembled genome for the *T. lucentensis* strain Mv1 was also deposited at GenBank under the same BioProject.

## Code Availability

Output from annotation tools, results files from the nf-core/rnaseq pipeline, and an R markdown script and html file for re-running R-analyses is provided at the figshare data repository: 10.6084/m9.figshare.29027924. The custom script “make_Mvib_annotations_table.py” for processing the functional annotation output into the overview presented in Data S6 is also available here.

## Acknowledgements

We thank Barry Stein for help with TEM imaging at the Electron Microscopy Center at Indiana University. We also thank Eduard Ocaña-Pallarès for advice and discussions This work was supported by the National Institutes of Health (R35GM138376) to J.P.G. The content of this paper is solely the responsibility of the authors and does not necessarily represent the official views of the National Institutes of Health. J.E.D is supported by an international postdoc grant from the Swedish Research Council (VR grant 734 2022-06250) and is a recipient of a non-stipendiary EMBO Long Term Fellowship (ALTF 740-735 2022). Work at the IR-T lab is co-funded by the European Union (ERC, MISSINGRELATIVES, 101097659). Views and opinions expressed are however those of the author(s) only and do not necessarily reflect those of the European Union or the European Research Council. Neither the European Union nor the granting authority can be held responsible for them. We also acknowledge support to Departament de Recerca i Universitats de la Generalitat de Catalunya (exp. 2021 SGR 00751). Computational analyses were enabled by resources in projects NAISS 2023/23-512, NAISS 2023/22-491, NAISS 2024/22-1422, and NAISS 2024/23-629 provided by the National Academic Infrastructure for Supercomputing in Sweden (NAISS) at UPPMAX, funded by the Swedish Research Council through grant agreement no. 2022-06725, and UPPMAX 2025/2-186 provided by Uppsala University at UPPMAX.

## Author contributions

Conceptualization, R.L., J.D., I.R.T., J.P.G.; methodology, R.L., J.D.; analysis, R.L., J.D., J.P.G.; investigation, R.L., J.D., K.K.; writing – original draft, R.L., J.D., J.P.G.; writing – review & editing, R.L., J.D., I.R.T., J.P.G.; visualization, R.L., J.D., J.P.G.; supervision, I.R.T., J.P.G.; funding acquisition, I.R.T., J.P.G.

## Competing interests

The authors declare no competing interests.

## Materials & Correspondence

Correspondence and requests for materials should be addressed to Joseph P. Gerdt & Iñaki Ruiz-Trillo.

## Extended Data Figures

**Extended Data Figure 1.**
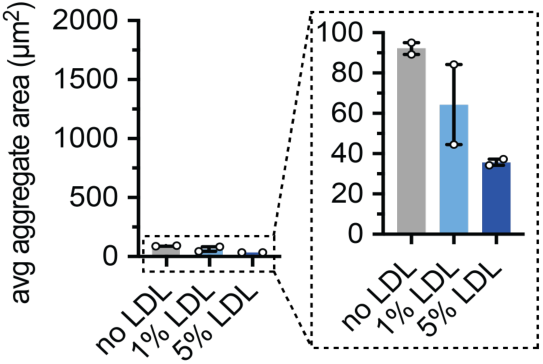
LDL does not induce aggregation in *M. vibrans*. Average aggregate area quantified for the xenic *M. vibrans* culture in the absence and presence of low density lipoproteins (LDLs, which induce aggregation in *C. owczarzaki*). Error bars represent the range of a biological duplicate. Individual replicates are displayed with circles.

**Extended Data Figure 2.**
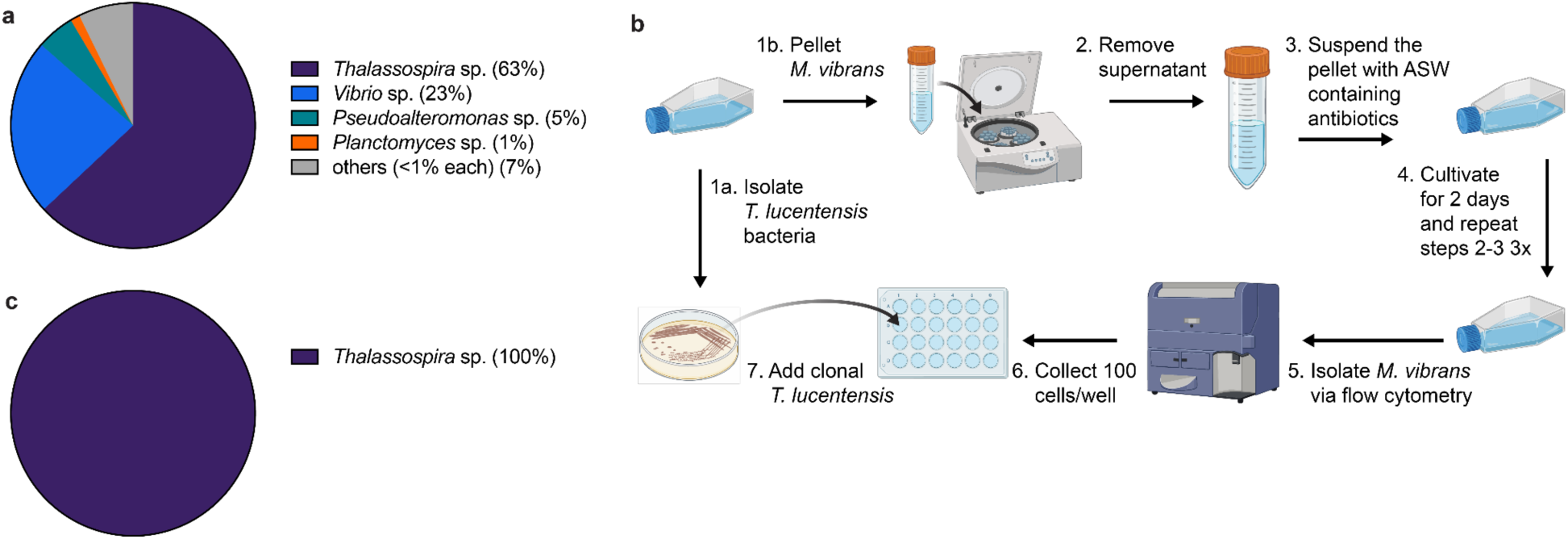
Generating a monoxenic *M. vibrans* culture. a) Results of 16S rRNA gene amplicon sequencing of the original xenic *M. vibrans* culture (Data S1). The four bacterial genera present at >1% of the population are indicated. b) Procedure to generate a monoxenic culture of *M. vibrans* with *T. lucentensis*. c) Result of 16S rRNA gene amplicon sequencing of the monoxenic *M. vibrans* culture. All reads were assigned to *Thalassospira*, originating from *T. lucentensis*. (Data S3)

**Extended Data Figure 3.**
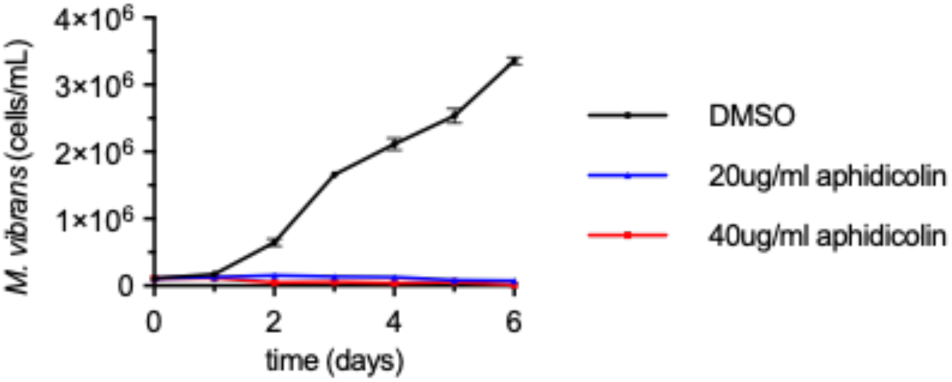
Validation of aphidicolin inhibition of *M. vibrans* replication. Growth curves of the monoxenic *M. vibrans* culture with aphidicolin (an inhibitor of replication) compared to a DMSO control. Scale bars represent standard error of the mean (SE) of a biological triplicate. The 20 µg/ml concentration was used in Fig 3c to arrest cell division.

**Extended Data Figure 4.**
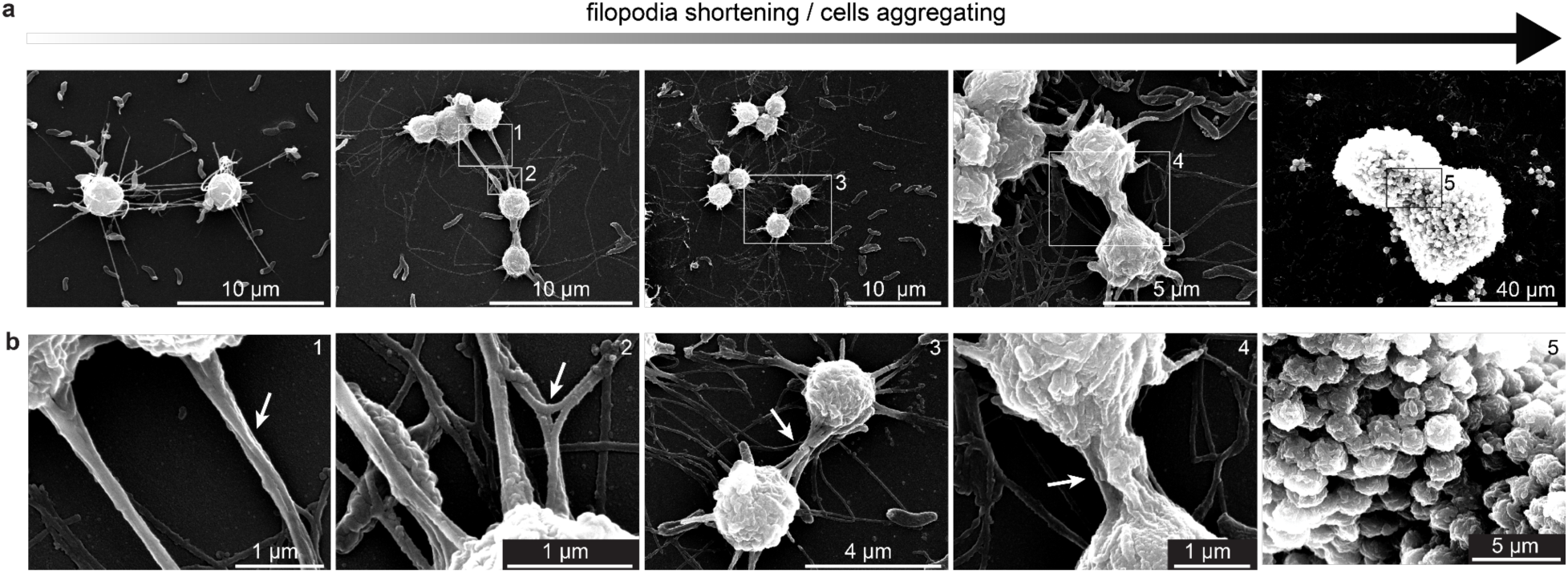
Model for the process of *M. vibrans* aggregation. a) Scanning electron microscopy images showing the putative process of aggregation from left to right. Filopodia of adjacent cells contact, tangle, and retract to form tight aggregates. b) Zoomed in images of the numbered white boxes. White arrows highlight entangled filopodia of adjacent cells.

**Extended Data Figure 5.**
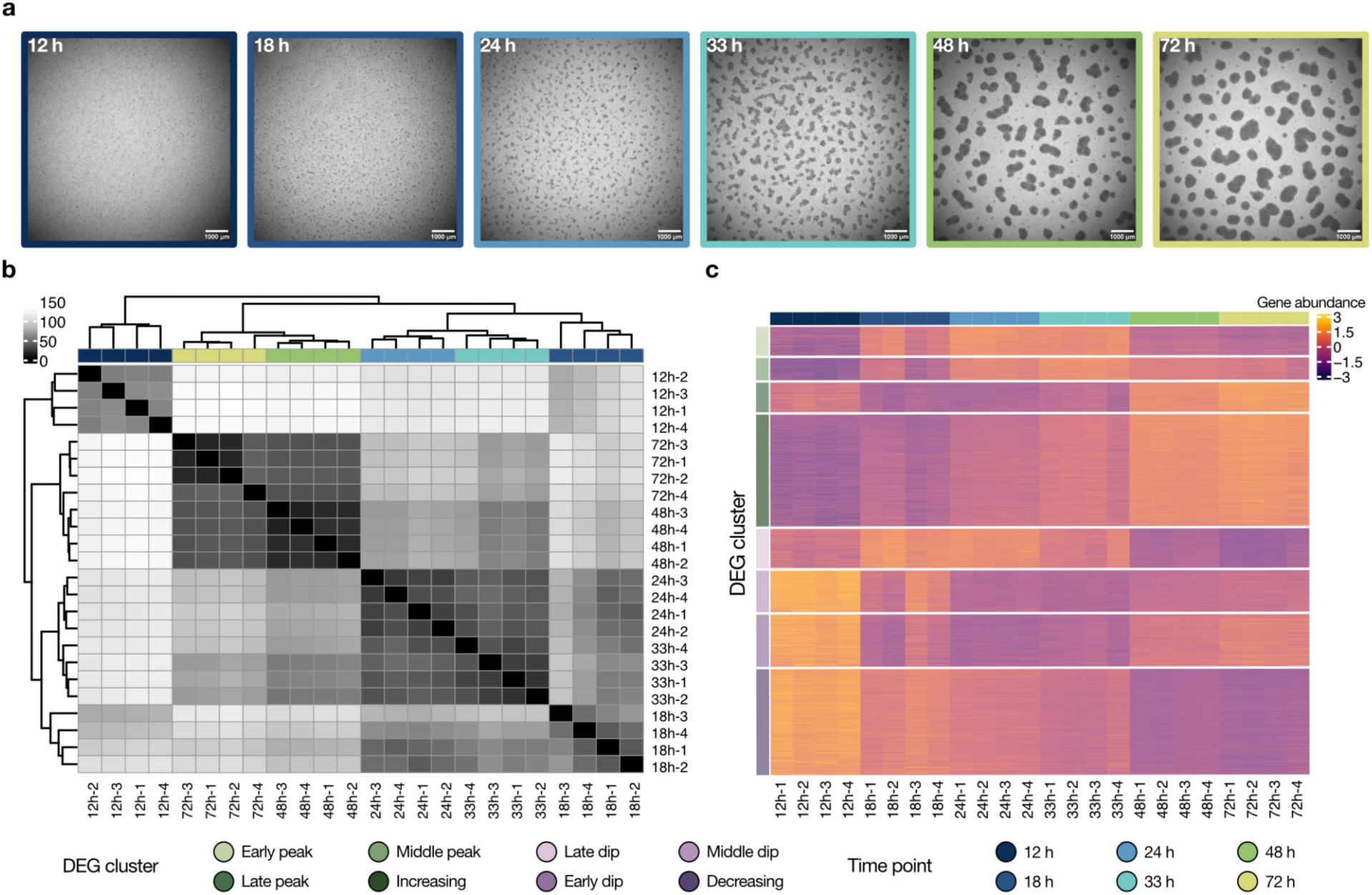
Time series of gene expression during the *M. vibrans* aggregation process. a) Representative images of aggregates across replicates at each time point where RNA was extracted for RNA-sequencing and downstream analysis of gene expression in the aggregation time series. b) Expression heatmap of sample-to-sample distances using the variance-stabilizing transformation of normalized gene counts from each of the four replicates sequenced at each of the six time points in the timeseries. c) Heatmap of gene abundances across the time series for all genes assigned to each DEG cluster. Gene abundances were centered and scaled using the variance-stabilizing transformation of normalized gene counts.

**Extended Data Figure 6.**
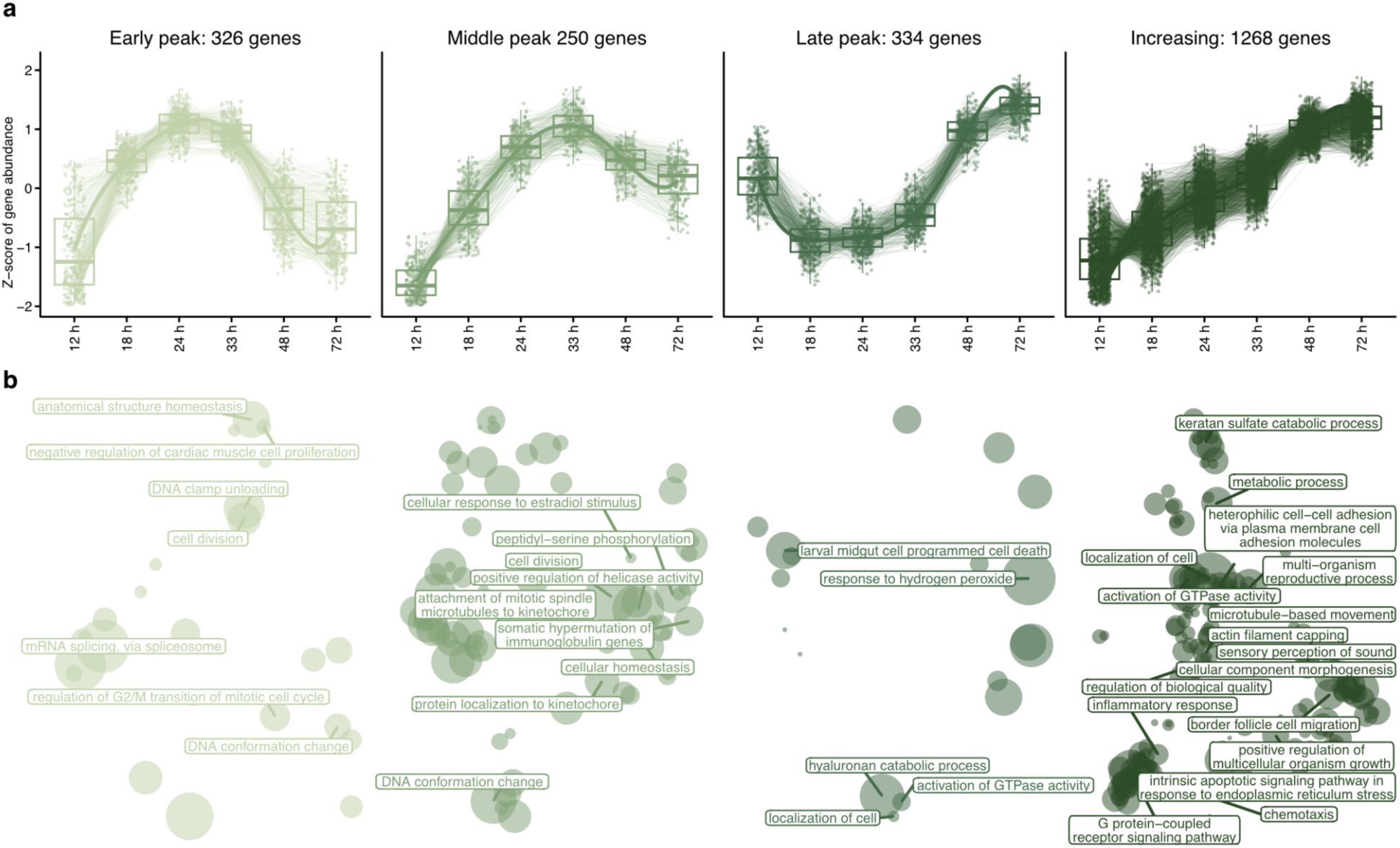
Upregulated groups of genes and enriched GO terms. a) Trends in abundance (Z-score scaled normalized gene counts) for genes with significant changes in expression across the time series assigned to the four DEG clusters, which are net upregulated. b) Scatterplots showing GO terms enriched in each differentially expressed gene (DEG) cluster. Distances between points in the plots indicate the similarity of terms, and the size of points represents their score. The names of the reduced set of GO terms based on semantic similarity and score are indicated.

**Extended Data Figure 7.**
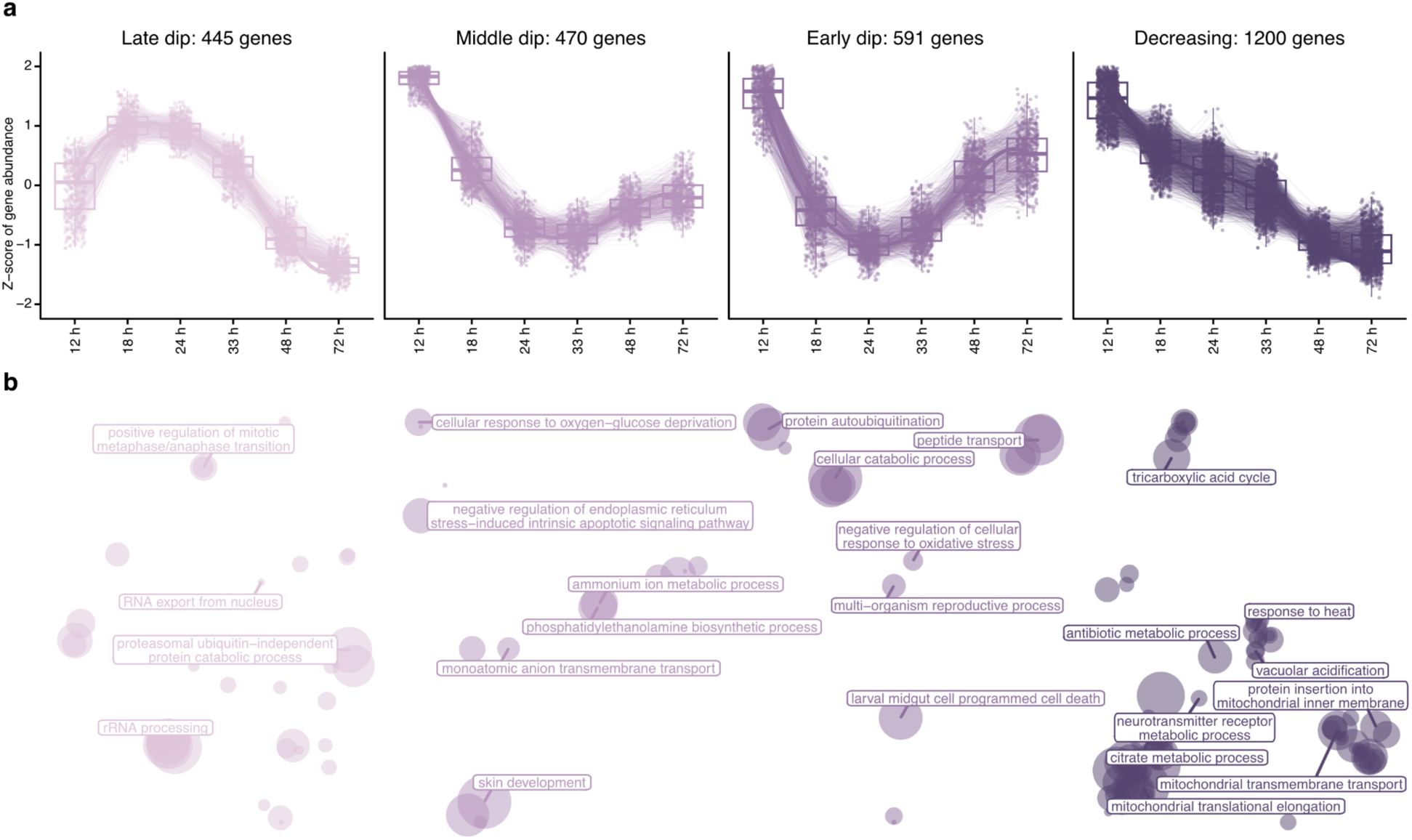
Downregulated groups of genes and enriched GO terms. a) Trends in abundance (Z-score scaled normalized gene counts) for genes with significant changes in expression across the time series assigned to the four DEG clusters, which are net downregulated. b) Scatterplots showing GO terms enriched in each differentially expressed gene (DEG) cluster. Distances between points in the plots indicate the similarity of terms, and the size of points represents their score. The names of the reduced set of GO terms based on semantic similarity and score are indicated.

**Extended Data Figure 8.**
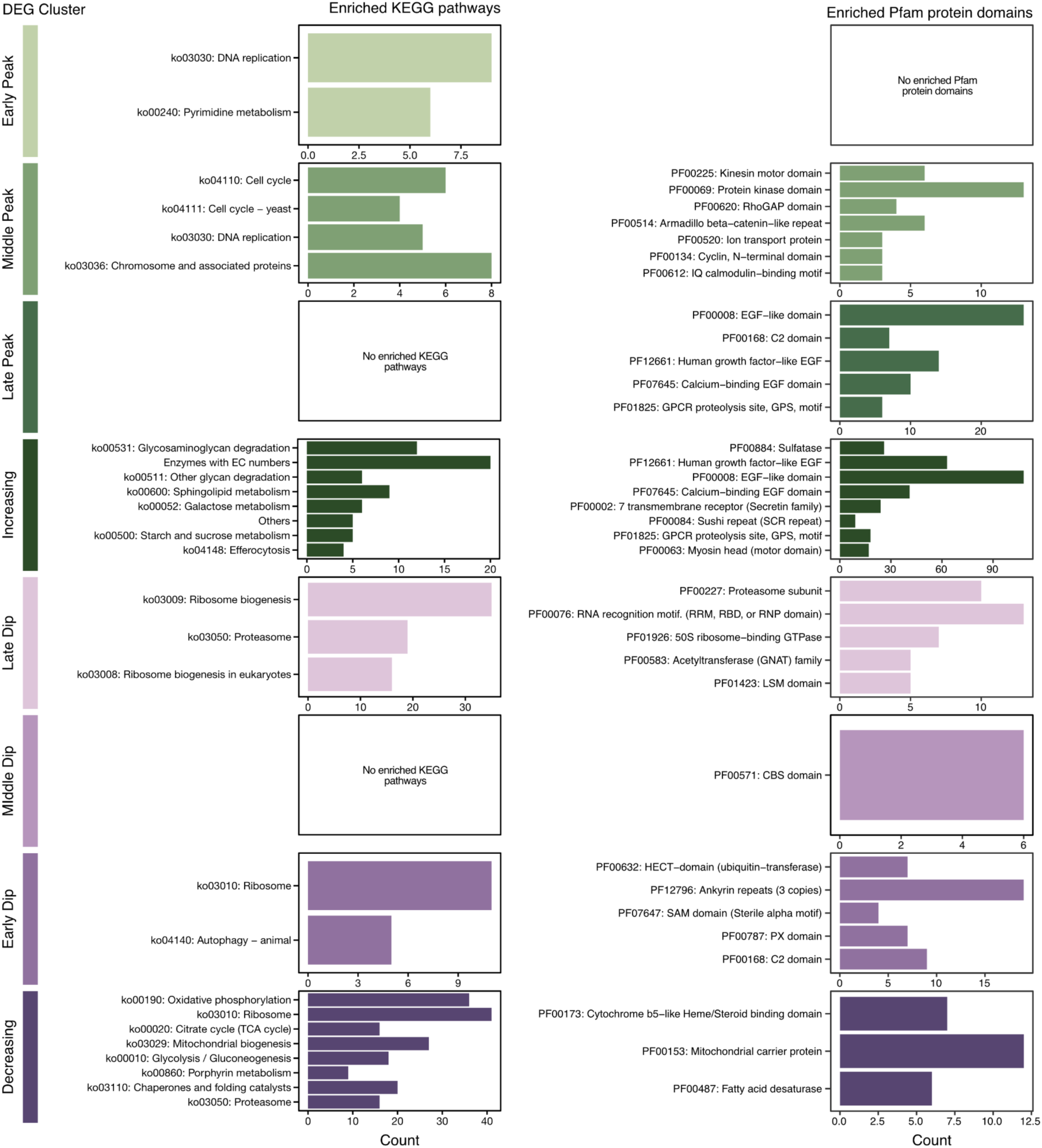
Pathways and protein domains expressed during *M. vibrans* aggregation. Overview of KEGG pathways and Pfam protein domains enriched in each differentially expressed gene (DEG) cluster, indicating which associated functions were expressed with varying patterns throughout the *M. vibrans* aggregation time series. For each plot, pathways and protein domains that meet the significance threshold (adjusted p-value ≤ 0.05) are ordered from lowest to highest adjusted p-value.

**Extended Data Figure 9.**
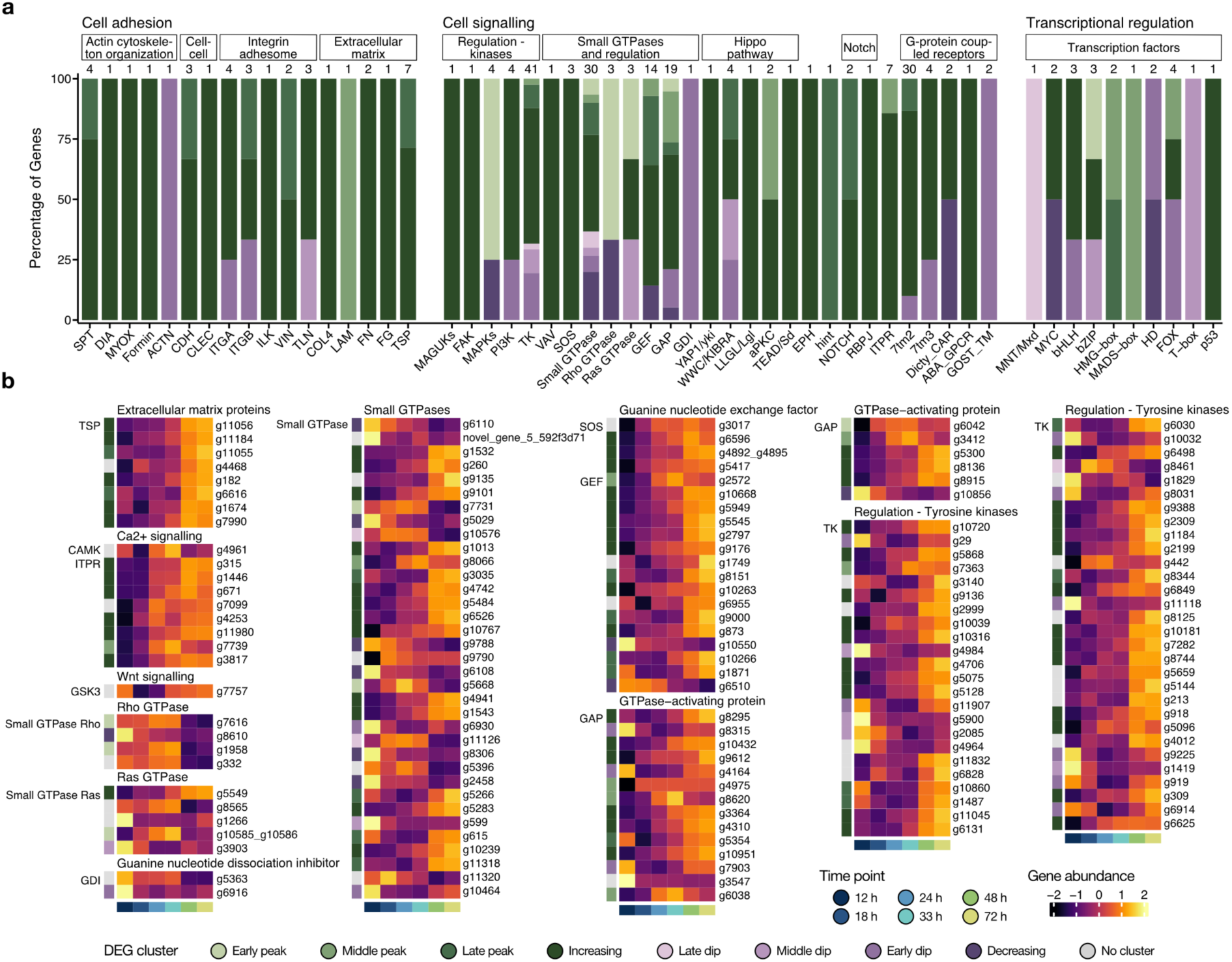
Diverse categories of *M. vibrans* multicellularity-related genes are differentially expressed during aggregation. a) Percentage of differentially expressed genes (DEGs) from each group of genes related to cell adhesion, cell signalling, and transcriptional regulation assigned to each DEG cluster. b) Heatmaps showing the expression patterns of additional genes involved in animal multicellularity that are significantly differentially expressed across the *M. vibrans* aggregation time series. The bottom row indicates the time point according to the legend and the left column DEG cluster assignment. Mean gene abundances were calculated across time point replicates and centered and scaled using the variance-stabilizing transformation of normalized gene counts. Thrombospondin; TSP, Ca^2+^/calmodulin-dependent protein kinase; CAMK, inositol 1,4,5_trisphosphate receptor; ITPR, glycogen synthase kinase 3; GSK3, guanine nucleotide dissociation inhibitor; GDI, Guanine nucleotide exchange factor; GEF, Son of Sevenless GEF; SOS, GTPase-activating protein; GAP, tyrosine kinases; TKs. See Fig. 5 for additional gene acronyms. See also Fig. S3 and Data S10.

**Extended Data Figure 10.**
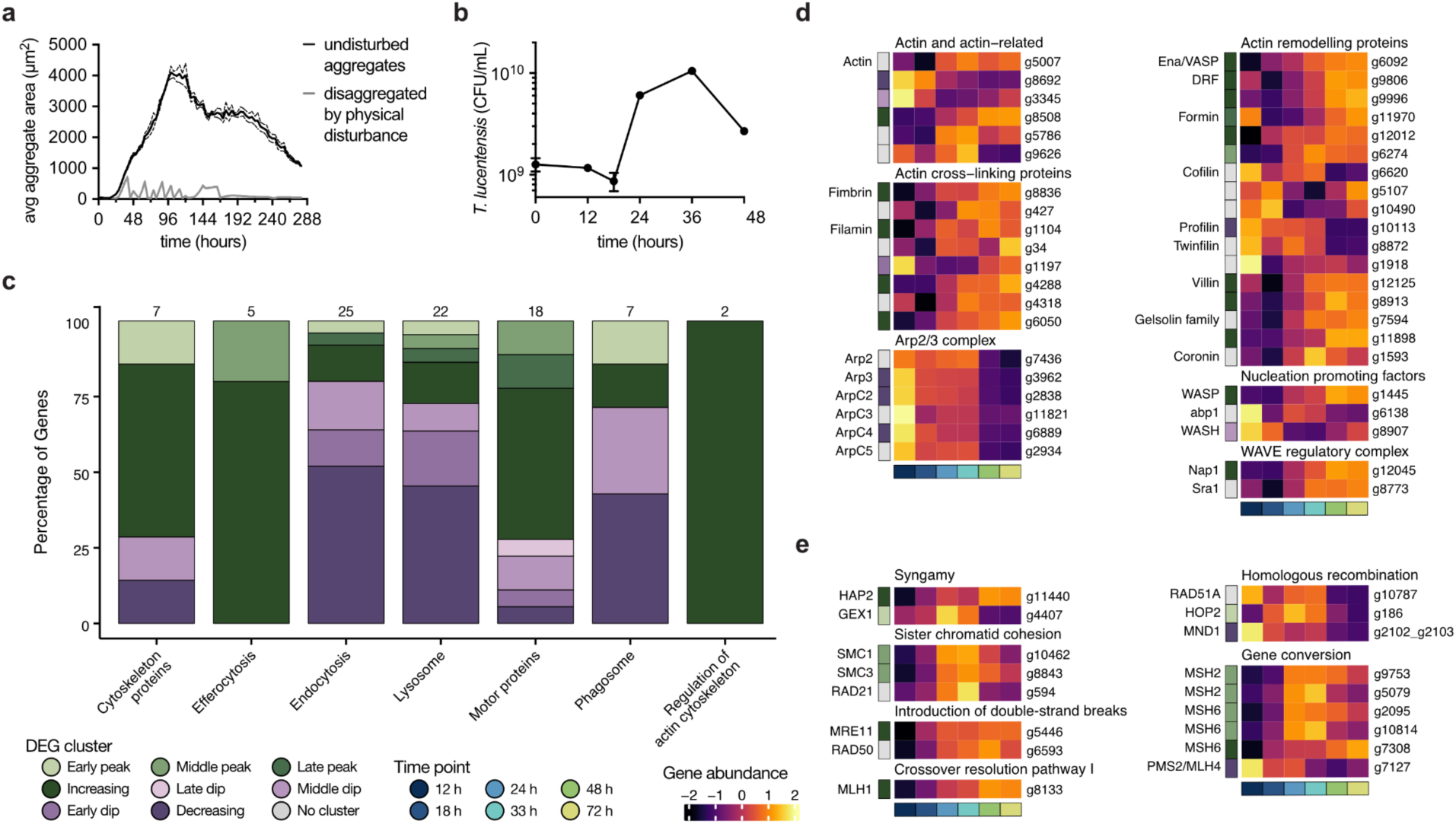
*M. vibrans* consumes bacteria, a) Average aggregate area quantified over time for *M. vibrans* cultures with/without physical disruption of aggregates. Dashed lines represent standard error of the mean (SE) of a biological triplicate. b) Colony-forming units of *T. lucentensis* food bacteria over time in the aggregating *M. vibrans* culture. Error bars represent standard error of the mean (SE) of a biological triplicate. c) Percentage of differentially expressed genes (DEGs) from phagocytosis-related KEGG pathways assigned to each DEG cluster. Heatmaps showing the expression patterns of d) phagocytosis and e) meiosis related genes significantly differentially expressed across the aggregation time series. The bottom row indicates the time point according to the legend and the left column DEG cluster assignment. Mean gene abundances were calculated across time point replicates and centered and scaled using the variance-stabilizing transformation of normalized gene counts. Diaphanous-related formin (DRF). See also Fig. S5 and Data S11.

## Notes

https://figshare.com/articles/dataset/dx_doi_org_10.6084/m9.figshare.29027924

### Competing Interest Statement

The authors have declared no competing interest.

