## Supplementary Information for "A close unicellular relative reveals aggregative multicellularity was key to the evolution of animals"

#### SUPPLEMENTARY DISCUSSION

##### Supplementary Discussion 1 – Enrichment analyses of differentially expressed gene clusters

To identify functions of importance for *M. vibrans* aggregation, we examined enriched Gene Ontology (GO) terms, Kyoto Encyclopedia of Genes and Genomes (KEGG) pathways, and Pfam protein domains across the eight differentially expressed gene (DEG) clusters. When considering the four DEG clusters composed of net upregulated genes (Fig. 4, Extended Data Figs. 6 and 8, Fig. S1, and Data S8–9), we found that functions related to DNA replication (e.g., ko03030), nucleotide metabolism (e.g., ko00240), mRNA splicing (e.g., GO:0000398), and the initiation of cell division (e.g., GO:0010389), were enriched among genes with peak expression early at 24 h in the *M. vibrans* aggregation process (early peak DEG cluster), and which subsequently drop in expression close to initial levels. Soon afterwards, functions related to the cell cycle (e.g., ko04110 and ko04111), kinetochore (eight GO terms), and mitosis (20 GO terms) reached peak gene expression at 33 h (middle peak DEG cluster) and remained elevated in comparison to initial expression levels. Unexpectedly, we found that eight GO terms related to meiosis were also enriched, suggesting that sexual reproduction may be occurring in *M. vibrans* aggregates. GO terms related to programmed cell death (GO:0035096) and cell localization and migration (five GO terms) were only later enriched among genes with peak expression in large mature aggregates at 72 h (late peak DEG cluster). The largest group of upregulated genes (constantly increasing DEG cluster) continued to increase in gene expression throughout the *M. vibrans* aggregation process. Strikingly, many of the enriched GO terms were associated with multicellularity and development, including cell adhesion (five GO terms) and cell signalling (35 GO terms), such as heterophilic cell-cell adhesion (GO:0007157), the G protein-coupled receptor (GPCR) signalling pathway (GO:0007186), regulation of Rho protein signal transduction (GO:0035023), MAPK cascade (GO:0000165), and positive regulation of hippo signalling (GO:0035332). Mirroring this, Pfam protein domains associated with both cell adhesion and signalling were also enriched and include epidermal growth factor-like domains (PF00008, PF07645, and PF12661), sushi repeats (PF00084), secretin 7 transmembrane receptors (PF00002), and G protein-coupled receptor proteolysis sites (PF01825). GO terms related to sexual reproduction, signalling involved in phagocytosis, negative regulation of immunity, and chemotaxis and cell localization are also enriched, in addition to KEGG pathways involving starch and glycan degradation (e.g., ko00500, ko00531, and ko00511).

Among the DEG clusters composed of net downregulated genes (Fig. 4, Extended Data Figs. 7–8, Fig. S2, and Data S8–9), we found that one group (late dip DEG cluster) initially underwent an increase in gene expression with a peak at 18 h before dropping in expression after 24 h. The functions enriched among these genes suggest an initial increase in transcription (i.e., RNA polymerase II preinitiation complex assembly, RNA export from nucleus, and mRNA processing), ribosome biogenesis (e.g., ko03008 and ko03009), translation (four GO terms), rRNA processing (ten GO terms), and the proteasome (i.e., ko03050), suggesting protein turnover. Another group of genes (early dip DEG cluster) underwent an initial decrease in gene expression at 24 h before increasing again, although not to initial expression levels. Here, enriched functions include autophagy (i.e., ko04140 and GO:0016236), programmed cell death (GO:0035096), and negative regulation of cellular response to oxidative stress (GO:1900408). Genes in the middle dip DEG cluster decreased in gene expression, reaching a dip at 33 h and remaining downregulated. Functions enriched in this group include stress responses such as oxygen-glucose deprivation (GO:0090650) and hypoxia (GO:1990910), and various catabolic and biosynthetic processes including tryptophan (GO:0006569), and amine (GO:0042402) catabolism, and tyrosine (GO:0006571), carnosine (GO:0035499), and phosphatidylethanolamine (GO:0006646) biosynthesis. The largest group of downregulated genes (constantly decreasing DEG cluster) continued to decrease in gene expression throughout the *M. vibrans* aggregation process and was enriched in many functions associated with central metabolism, including glycolysis/gluconeogenesis (ko00010), the TCA cycle (ko00020), and oxidative phosphorylation (ko00190). Mitochondrial biogenesis (ko03029) and mitochondrial import and electron transport (17 GO terms) were also particularly prevalent among enriched functions.

#### Supplementary Discussion 2 – Animal multicellularity-related genes are expressed during *M. vibrans* aggregation

Stable and organized adhesion between neighboring cells is a critical requirement for the evolution of complex multicellularity. Extant animals achieve this through the deployment of genes for structural remodeling and cellular scaffolding, such as those involved in actin cytoskeleton organization, producing extracellular matrix (ECM) to form a basement membrane for cell and tissue support, and protein complexes that mediate cell-ECM and cell-cell adhesion. An extensive repertoire of these cell adhesion genes is also found encoded in the genomes of unicellular holozoans, although in many cases we do not know their function in this context.[1-15] We found that the large majority of *M. vibrans* cell adhesion DEGs were upregulated (Fig. 5, Extended Data Fig. 9, Supplementary Fig. 3, and Data S10). These included the actin filament bundling genes spectrin (SPT), fascin (FSCN), and diaphanous-related formin (DIA) gained in the urfilozoan ancestor, and myosin X (MYOX) gained in the urholozoan ancestor. SPT is a cytoskeletal protein necessary for resisting mechanical stress and epithelia formation in animals, it has previously been described in choanoflagellates and protein domains are also found in *C. owczarzaki*.[1,10,16,17] FSCN is involved in animal filopodia and microvilli actin cytoskeleton organization, with the *S. rosetta* homolog localizing to analogous structures and *C. owczarzaki* homologs expressed in adherent cells.[4] DIA functions in actin filament nucleation, and MYOX is essential for filopodia formation in animals.[5,18] Diverse homologs of ECM genes were also upregulated during *M. vibrans* aggregation, including a previously identified putative alpha type IV collagen (COL4; PF01413), which is the only case found thus far outside Metazoa (Fig. 5b–d).[12] In addition, genes encoding laminin G (LAM G, PF02210), fibronectin type II and III (FN I; PF00040, and FN III; PF00041), and the fibrinogen C-terminal (FG C; PF00147) protein domains were upregulated, having a peak in expression either during aggregate formation or in large mature aggregates (Fig. 5c). We also found that a large number of genes encoding thrombospondin (TSP) type 1 domains (PF00090), inferred to have a urholozoan origin, were upregulated (Extended Data Fig. 9). However, no putative thrombospondin genes encoded the C-terminal domain (PF05735) found in some choanoflagellates.[13] Genes encoding several of these ECM protein domains have previously been predicted to be secreted by the choanoflagellate *Monosiga brevicollis* and *C. owczarzaki*.[9] In terms of cell-ECM adhesion, we identified several dystroglycan complex gene homologs in *M. vibrans*, including a putative syntrophin, dysbindin, dystrophin, and sarcoglycan (Data S10). However, none were differentially expressed across the time series. On the other hand, homologs of most integrin adhesome genes were upregulated. The integrin adhesome plays a central role in cell-ECM adhesion in animals and has ancient origins in the last common ancestor of Amorphea.[3,19,20] Homologs of integrin alpha (ITGA), integrin beta (ITGB), talin (TLN), paxillin (PXN), and integrin-linked kinase (ILK) were all upregulated, although PINCH and parvin (PARV) were not significant DEGs (Fig. 5c and Data S10). In addition, we found that homologs of tyrosine-protein kinase c-Src (SRC), focal adhesion kinase (FAK), and C-terminal Src kinase (CSK) were all upregulated, which function in integrin-mediated signalling and are holozoan-specific.[11] The alpha-catenin/vinculin family (VIN; PF01044) includes both vinculin and alpha-catenin, which function in the integrin adhesome and cell-cell adhesion, respectively and cannot be distinguished in non-metazoans.[7] Mirroring *C. owczarzaki*, we found that *M. vibrans* also encodes three VIN homologs, two of which are upregulated during aggregation. *M. vibrans* encodes several additional cell-cell adhesion homologs, including putative cadherin (CDH), C-type lectin (CLEC), and ezrin-radixin-moesin (ERM) family genes. CDHs are the major components of adherens junctions, which mediate cell-cell adhesion in animals. CDH has a urfilozoan origin and *C. owczarzaki* encodes only one homolog, while *M. vibrans* encodes seven (eight with PF00028, Data S10), on the same order as ctenophores, placozoans, and some sponges, yet fewer than the 23 and 29 found in *M. brevicollis* and *S. rosetta*.[2,6,7] Three *M. vibrans* CDH homologs are DEGs and all are upregulated, including two genes (g1326 and g10477) that increased over 15 times in gene expression over the time series (Fig. 5b–d and Data S10). Canonical CLEC domains (PF00059) have a patchy distribution in Holozoa and were previously reported to be absent in Filasterea.[15] However, we found that *M. vibrans* encodes two genes with the domain, and a third with a different CLEC domain (PF19439), one of the former (g5462) is a DEG and upregulated during aggregation. The ERM gene family is holozoan-specific and includes cytoskeletal proteins involved in forming cell-cell contacts and merlin, a gene in the Hippo signalling pathway, which cannot be distinguished between unicellular Holozoa.[12,14] Of the four *M. vibrans* ERM homologs, one was a DEG with a peak in gene expression at 33 h (Fig. 5c and Data S10).

Cell signalling mechanisms involved in intracellular signal transduction, immunity, environmental signal/response pathways, and cell-cell communication are also critical for evolving complex multicellularity. Animal multicellular development is mediated by a small set of specific cell signalling pathways and receptors (*e.g.*, Hippo, Hedgehog, Wnt, TGF-beta, Notch/Delta, toll-like receptors (TLRs), teneurins, membrane-associated guanylate kinases (MAGUKs), nuclear hormone receptors (NR), Ephrin-Eph, and JAK-STAT) alongside eukaryote-wide cell signalling mechanisms that have undergone gene expansions and/or subfunctionalization (e.g., tyrosine kinases (TKs) and receptor tyrosine kinases (RTKs), mitogen-activated protein kinases (MAPKs), Phosphatidylinositol-3-kinases (PI3Ks), Small GTPases and regulators, $Ca^{2+}$ signalling, and G-protein coupled receptors (GPCRs)).[21-27] Homologs of genes from several specific cell signalling pathways and receptors have now been identified outside of Metazoa, primarily in unicellular Holozoa, including Hippo, Hedgehog, Ephrin-Eph, Notch/Delta, teneurins, MAGUKs, TLRs, and teneurins.[1,6,16,24,28-32] We identified many cell signalling gene homologs in the *M. vibrans* genome, with the majority of DEGs upregulated across the aggregation time series (Fig. 5, Extended Data Fig. 9, Supplementary Fig. 3, and Data S10). In terms of signal transduction, we found 101 predicted TKs (PF07714), a high proportion of which were upregulated (Extended Data Fig. 9 and Data S10). Further analyses are necessary to determine which are RTKs, but 103 TKs are also found in *C. owczarzaki*, most of which were RTKs, and several RTKs were previously described in *M. vibrans* through a PCR-based survey.[22] In addition, *M. vibrans* encodes seven putative MAPKs, three of which had an early peak in gene expression (Fig. 5c and Data S10). We also identified five PI3Ks, with three upregulated, while the downstream mTOR decreased in expression in large mature aggregates. In animals, MAGUKs regulate the formation of cell-cell adhesions, once thought to be Metazoa-specific they are also found in choanoflagellates and *C. owczarzaki*.[33] We identified four MAGUK homologs, one which was upregulated, a second which peaked in expression during aggregation, and a third, a putative disc large MAGUK (DLG), that dropped in expression in large mature aggregates. Small GTPases and their regulators are key signalling components in central cellular processes in eukaryotes, where they act as molecular switches in response to various signals including from receptors.[34] We annotated 81 Small GTPases (PF00071), of which eight are putative Ras GTPases (IPR020849) and ten putative Rho GTPases (IPR003578), 28 guanine nucleotide exchange factors (RhoGEF, PF00621) that activate GTPases, 31 GTPase-activating proteins (RhoGAP, PF00620) that negatively regulate GTPases, and three guanine nucleotide dissociation inhibitors (RhoGDI, PF02115) that sequester GTPases (Data S10). Expression patterns among the small GTPases were more variable, but the majority of RhoGEFs and RhoGAPs were upregulated, while RhoGDIs were downregulated (Extended Data Fig. 9). In particular, we found that the RhoGEF VAV, which is filozoan-specific[35] was upregulated (Fig. 5b–c). We also found that growth factor receptor-bound protein 2 (GRB2), a signalling adaptor that links RTKs to ras signalling, was upregulated. Regarding specific signalling pathways, we found homologs of both Ca2+/calmodulin-dependent protein kinase (CAMK) and inositol 1,4,5-trisphosphate receptors (ITPR/IP3Rs), both involved in $Ca^{2+}$ signalling, with CAMK having a middle peak in gene expression, and most IP3Rs upregulated (Extended Data Fig. 9). In animals, the Hippo pathway regulates cell proliferation, apoptosis, and tissue size, with many components having pre-metazoan origins.[30] In the *M. vibrans* genome we identified homologs of Yes-associated protein 1/Yorkie (YAP1/yki), WW and C2 domain containing/kidney and brain expressed protein (WWC/KIBRA), Lethal (2) giant larvae (LLGL/Lgl), protein kinase C (aPKC), Large tumor suppressor kinase/Warts (LATS/Wts), TEA domain transcription factor/scalloped (TEAD/Sd), Macrophage stimulating 1/Hippo (MST1/Hpo), and MOB kinase activator/mob as tumor suppressor (MOB/Mats) (Data S10). Of these, homologs of all holozoan and filozoan specific-genes were upregulated (*i.e.*, YAP1/yki, WWC/KIBRA, LLGL/Lgl, and aPKC) alongside TEAD/Sd, while MOB/Mats dropped in expression in large mature aggregates (Figure 5b–d). In *C. owczarzaki*, YAP1/yki regulates the shape of multicellular aggregates,[36] and its upregulation during *M. vibrans* aggregation could suggest a similar role. Most components of the Hedgehog pathway are animal-specific where the pathway is central in regulating cell differentiation.[24] However, choanoflagellates encode genes with either hedge (*i.e.*, hedgling) or hint/hog domains that together are found in hedgehog (HH/hh), and the hedgehog receptor patched (PTCH), homologs of which are also found in other eukaryotes.[1,6,28,37,38] Unexpectedly, we identified three *M. vibrans* genes that encoded a hint/hog domain (PF01079) and were reciprocal best hits (RBH) to human hedgehog homologs, alongside a putative PTCH gene (Data S10). One of the hint homologs was upregulated, alongside PTCH which underwent a sharp increase in expression at 18 h (Fig. 5c). With the exception of Glycogen synthase kinase 3 (GSK-3) from the Wnt pathway, and several TGF-β related protein domains found in choanoflagellates, the Wnt, TGF-β, NR, and JAK pathways and receptors remain animal-specific, with no homologs found in *M. vibrans* (Data S10).[24,31] In terms of signalling receptors, we found that *M. vibrans* encodes an extensive repertoire of GPCRs: one Rhodopsin/Class A (7tm1, PF00001), 51 Adhesion/Secretin/Class B (7tm2, PF00002), six Glutamate/Class C (7tm3, PF00003), two cAMP/ClassE (Dicty_CAR, PF05462), one Abscicic acid receptor (ABA_GPCR, PF12430), and seven GOST seven transmembrane (GOST_TM, PF06814) genes (Data S10). *M. vibrans* encodes an exceptional number of 7tm2, on the same order as metazoans (de Mendoza 2014), and the vast majority that are differentially expressed are upregulated (Fig. 5c). Most 7tm3 genes were also upregulated, alongside ABA_GPCR and one Dicty_CAR, while 7tm1 and GOST_TM were downregulated. We also identified one putative eph receptor (EPH; g733) from the Ephrin-EPH pathway, which in animals is critical for morphogenesis and filozoan-specific,[32] which increased over 10x in gene expression over the course of the aggregation time series (Figure 5b–d). In addition, we identified three putative Notch receptor (NOTCH) gene homologs, which is filozoan-specific,[31] all of which were upregulated alongside recombining binding protein suppressor of hairless (RBPJ), a transcription factor (TF) in the pathway.

Transcriptional regulation through the activity of TFs is crucial for coordinating gene deployment in animal development. The TF repertoires of complex multicellular lineages are likewise more complex, although they were assembled in a stepwise manner and many key animal TF families have pre-metazoan origins.[3,39-43] In the *M. vibrans* genome we identified a large repertoire of TFs important in Metazoa: eight basic helix loop helix (bHLH) including Myc-Max network homologs (MAX, MNT/Mxd, MYC), four basic leucine zipper (bZIP), seven high mobility group (HMG) box including a putative SRY-related (SOX) gene, two MADS-box including a putative Myocyte-specific enhancer factor 2 (MEF2) gene, six homeodomain (HD) including a putative LIM gene, two TALE-related HD, eight forkhead (FOX), one CP2 that is grainyhead-like (GRH), one Churchill, two T-box including Brachyury, three Runt-related (RUNX), one p53, one Rel homology DNA-binding (RHD), and two regulatory factor X (RFX) (Data S10). In contrast to cell adhesion and signalling, a nearly equal number of transcriptional regulation genes that were differentially expressed were net upregulated and downregulated across the *M. vibrans* aggregation time series (Fig. 5a). The individual expression profiles of these TFs were diverse, with the overarching patterns of peaking in expression prior to the initiation of aggregate formation, sharply increasing in expression at the onset of aggregation, peaking in expression in small or large aggregates, or constantly increasing in expression (Fig. 5, Extended Data Fig. 9, Supplementary Fig. 3, and Data S10). These patterns likely reflect the functions of different TFs in regulating the expression of specific gene sets both prior to and throughout the *M. vibrans* aggregation process. A number of these TFs are specific to Opisthokonta, Holozoa, or Filozoa and show strong expression patterns (Fig. 5b–c). For example, the differentially expressed T-box TF, which are Opisthokonta-specific and regulate metazoan development,[3] had a sharp peak in expression at 12 h, prior to the initiation of aggregation. The putative SOX on the other hand had a sharp peak in expression at 33 h, prior to the formation of large mature aggregates. SOX TFs are filozoan-specific, and choanoflagellate SOX can induce the formation of mouse stem cells from somatic cells, mirroring the function of SOX in animals (Gao 2024). Both RUNX and p53, which are Holozoa-specific,[3,41] sharply increased in expression at the onset of aggregation (Fig. 5b–d). RHD on the other hand, which emerged in Opisthokonta and includes TFs regulating immunity, development, and differentiation in animals,[42] reached peak expression in large mature aggregates.

**Supplementary Discussion 3 – Phagocytosis- and meiosis-related genes are expressed during *M. vibrans* aggregation**

There are few highly conserved phagocytosis components found across eukaryotes that do not have prokaryotic homologs and these genes can have diverse functions. However, phagocytosis depends on actin cytoskeleton remodelling and branched filament formation through the action of Arp2/3.[44] Unicellular holozoans have also previously been shown to encode diverse proteins associated with actin-based cellular protrusions necessary for phagocytosis.[18] We thus searched for homologs of such phagocytosis genes in the *M. vibrans* genome and investigated their individual expression profiles, in addition to investigating the patterns of DEGs in related KEGG pathways (Extended Data Fig. 5, Fig. S5, and Data S11). We found that different actin homologs were expressed both early and later during aggregation. In terms of the Arp2/3 complex, all components were highly upregulated until just prior to the formation of small aggregates before dropping in expression. Some actin remodelling proteins were upregulated initially (i.e., cofilin, profilin, and twinfilin) before a switch where the majority of actin remodelling proteins become upregulated as large mature aggregates formed (i.e., Ena/VASO, diaphanous-related formins, formins, villin, gelsolin family, and coronin), alongside the upregulation of genes involved in the WAVE regulatory complex. These results suggest an initial burst in actin polymerization prior to aggregate formation later followed by actin remodelling in large mature aggregates. The KEGG pathway DEG patterns were consistent with this as endocytosis, phagosome, and lysosome pathways had a higher proportion of DEGs that were net downregulated, and thus related genes were highly expressed prior to aggregate formation. While cytoskeleton proteins, efferocytosis, motor proteins, and regulation of actin cytoskeleton pathways had a higher proportion of DEGs that were net upregulated, suggesting an increase in expression after aggregate formation. Together these results are consistent with our experimental findings that bacterial feeding through phagocytosis is occurring in *M. vibrans* aggregates. Consistent with our results showing increased growth in *M vibrans* aggregates (Fig. 6d), *S. rosetta* colonial multicellular rosettes have previously been shown to increase feeding efficiency through higher rates of food vacuole formation compared to unicellular cells.[45]

Sexual reproduction has previously been demonstrated in the choanoflagellate *S. rosetta*, which undergoes meiosis in response to stress from nutrient limitation[46] or bacterial chondroitinases, such as produced by *Vibrio fischeri*.[47] Genes involved in meiosis are found across diverse eukaryotes including all sequenced lineages of unicellular holozoans.[48] HAP2 and GEX1 in particular are meiosis-specific as they are essential for gamete fusion and karyogamy, respectively, and have been used as meiosis marker genes, including in unicellular holozoans.[49-51] In particular, an extensive complement of genes from the meiosis gene toolkit[52] has previously been detected in both *M. brevicollis*[53] and *C. owczarzaki.*[16] Given the enrichment of GO terms related to meiosis during *M. vibrans* aggregation, we searched for homologs of meiosis-related genes and investigated their individual expression profiles (Fig. 6h, Extended Data Fig. 10, Fig. S5, and Data S11). We focused on genes previously investigated in a study on the distribution of meiosis across Amoebozoa.[54] We found that *M. vibrans* encodes nearly a full complement of meiosis-associated genes, including the meiosis-specific GEX1 and HAP2, both of which are highly upregulated in small aggregates and large mature aggregates, respectively. Genes related to homologous recombination were also upregulated early during aggregate formation and became downregulated in large mature aggregates. While genes related to sister chromatid cohesion and gene conversion peaked in expression in small aggregates. Genes related to the introduction of double-strand breaks and the crossover resolution pathway I, on the other hand, continued to increase in expression in large mature aggregates. Overall, our results are suggestive of meiosis occurring in M. vibrans aggregates. As an aside, we also found that *M. vibrans* encodes a putative chondroitin lyase (g557) as it has the same predicted glycosaminoglycan (GAG) lyase domains (PF08124, PF02278, and PF02884) as EroS, which is the *V. fischeri* mating inducer.[47] The gene increases over four times in expression over *M. vibrans* aggregation and is a top hit to the *S. rosetta* sequence XP_004988474.1, which is also a chondroitin lyase.

### SUPPLEMENTARY FIGURES

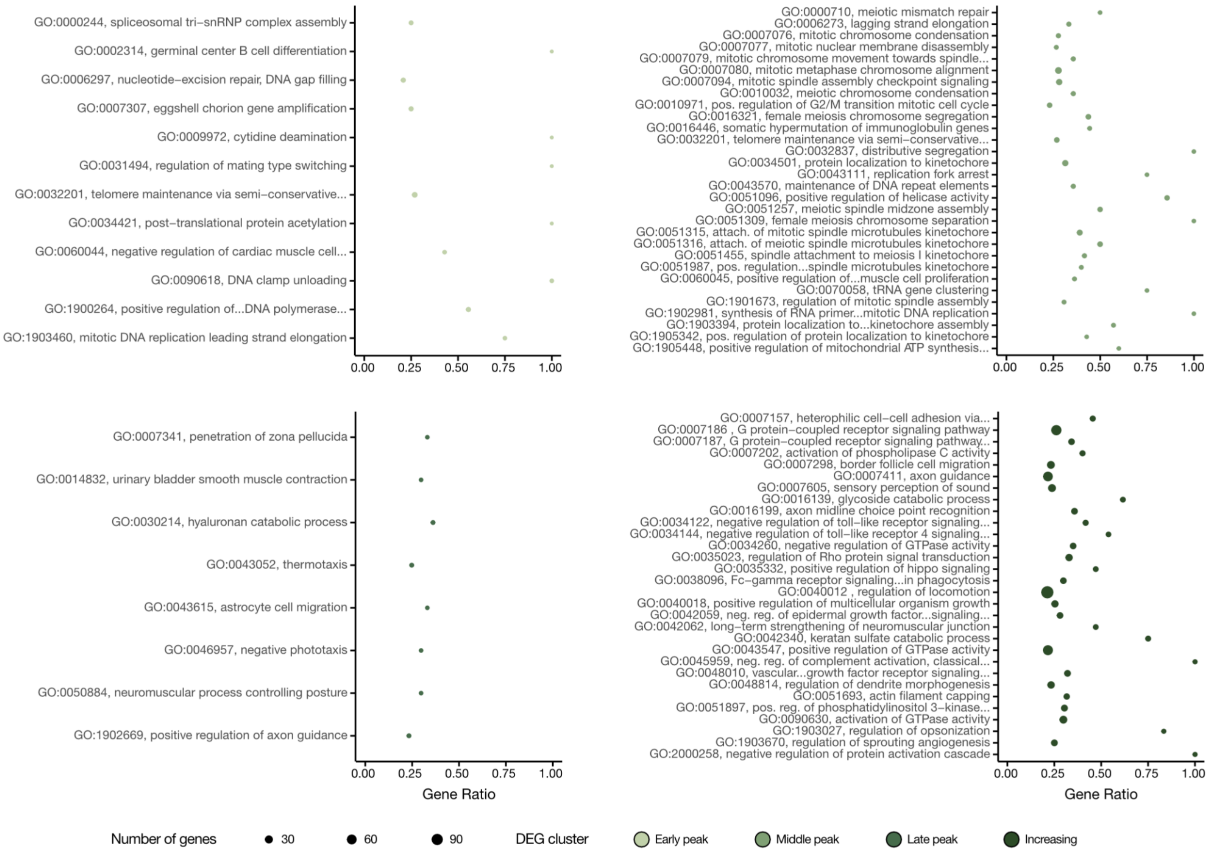


**Figure S1.** Dotplot overview of the top 30 enriched GO terms with the lowest weighted Fisher p-values ($p \leq 0.001$) across each of the four net upregulated differentially expressed gene (DEG) clusters. Gene ratio indicates the proportion of genes assigned to each GO term that are cluster DEGs, terms with less than 0.2 were excluded. Dot size indicates the number of cluster DEGs assigned to the corresponding GO term. Colours indicate the DEG cluster. See Data S9 for all enriched GO terms and complete names.

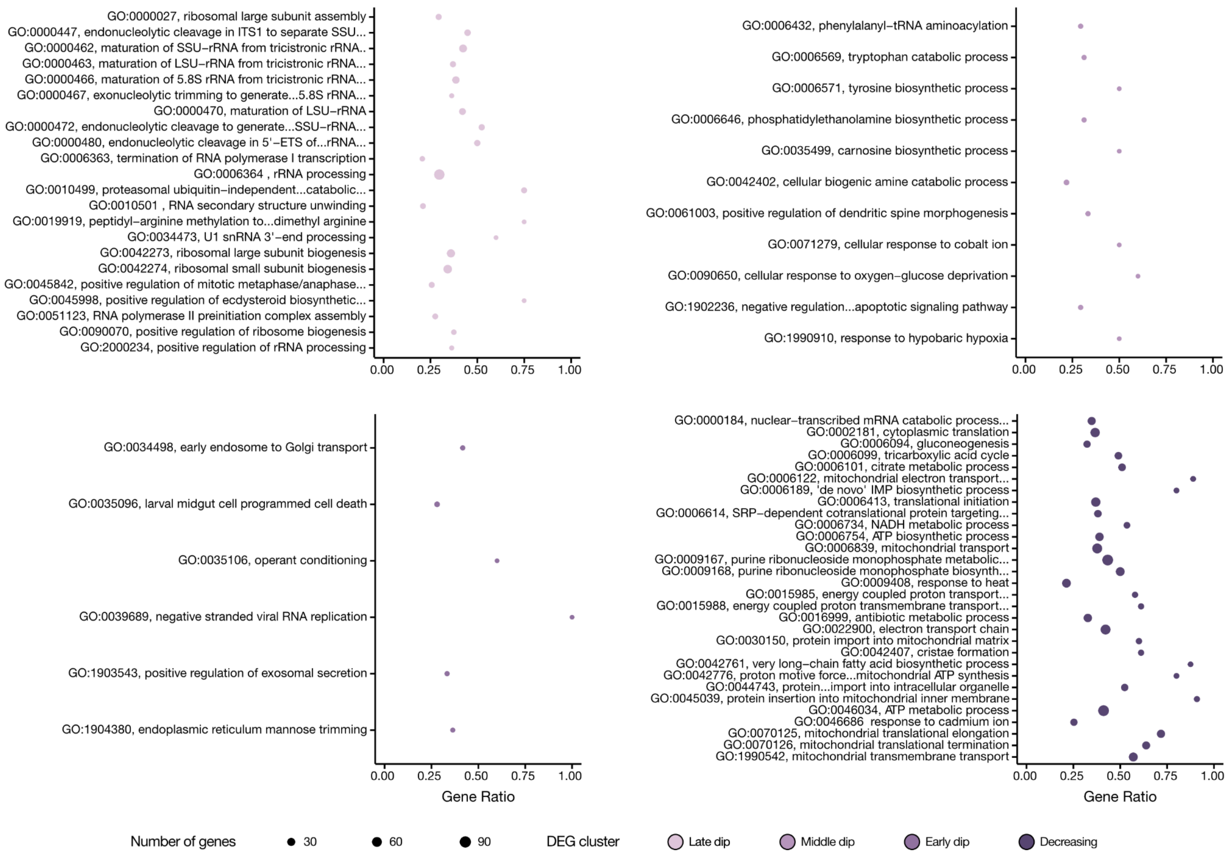


**Figure S2.** Dotplot overview of the top 30 enriched GO terms with the lowest weighted Fisher p-values ($p \leq 0.001$) across each of the four net downregulated differentially expressed gene (DEG) clusters. Gene ratio indicates the proportion of genes assigned to each GO term that are cluster DEGs, terms with less than 0.2 were excluded. Dot size indicates the number of cluster DEGs assigned to the corresponding GO term. Colours indicate the DEG cluster. See Data S9 for all enriched GO terms and complete names.

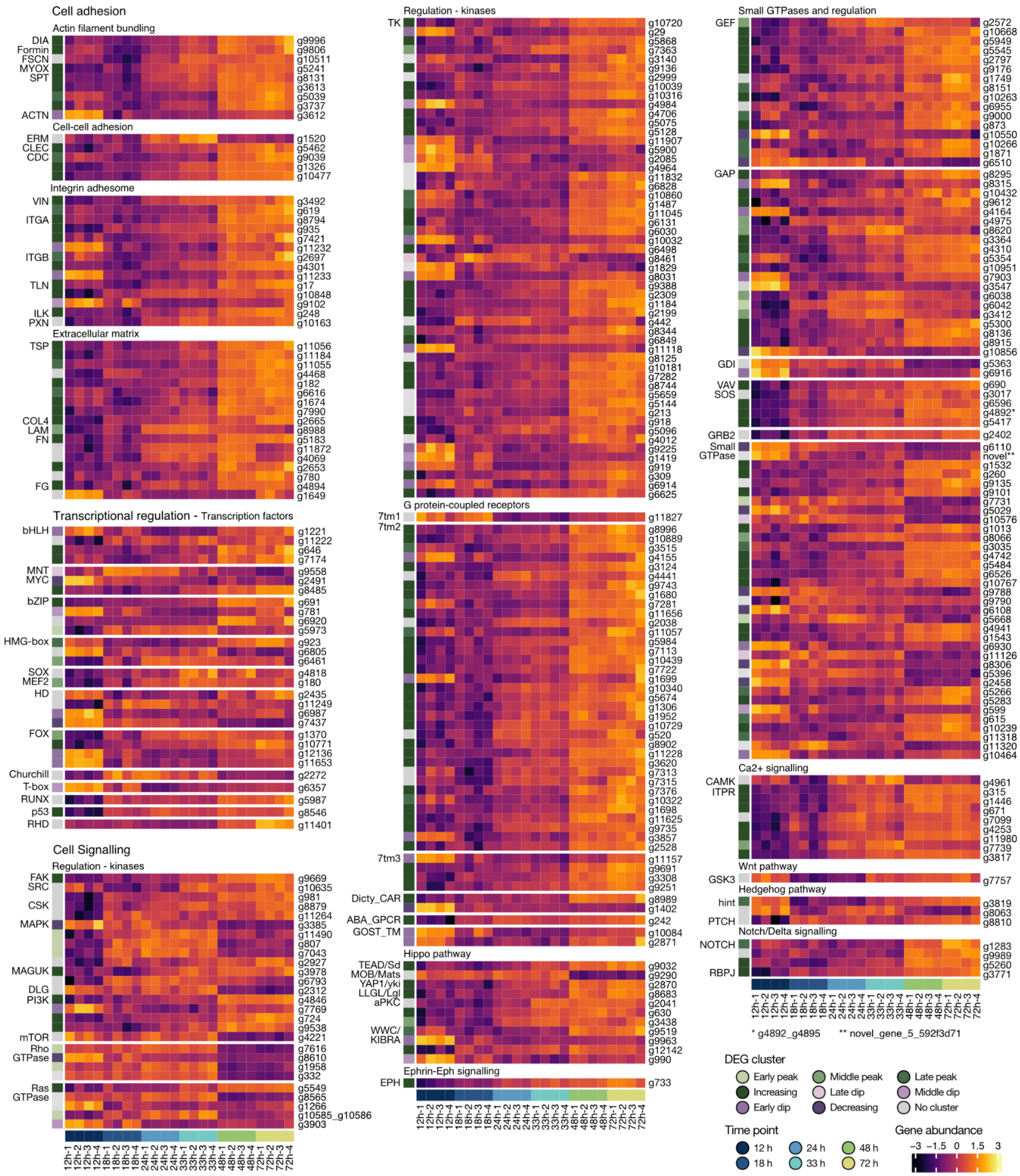

**Figure S3.** Heatmaps showing the expression patterns of genes involved in animal multicellularity that were significantly differentially expressed across the *M. vibrans* aggregation time series. The bottom row indicates the time point according to the legend and the left column DEG cluster assignment. Gene abundances for replicates at each time point were centered and scaled using the variance-stabilizing transformation of normalized gene counts. See Fig. 5 and Extended Data Fig. 9 for gene acronyms.

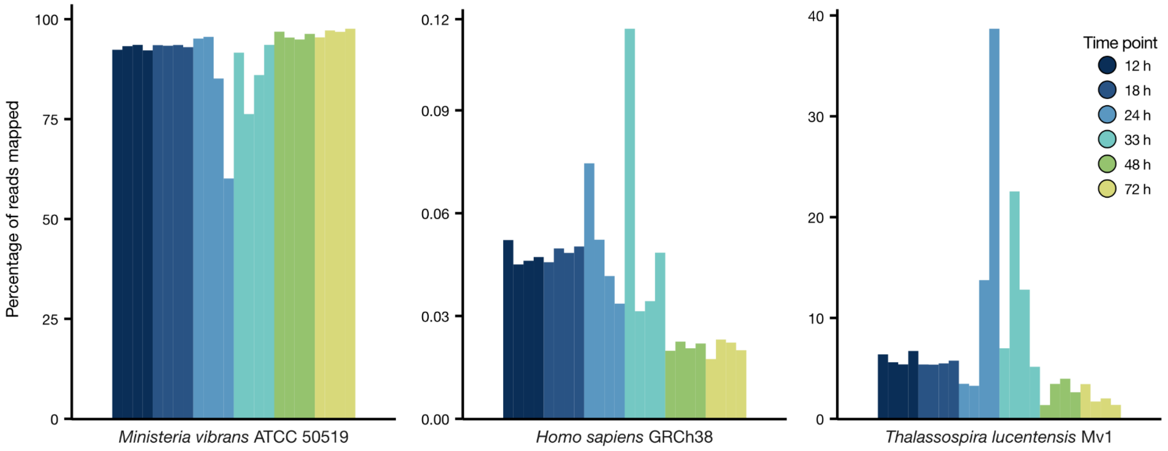


**Figure S4.** Percentage of RNA-sequencing reads from each of the four replicates at each time point in the aggregation time series mapped against the *M. vibrans* ATCC 50519, *Homo sapiens* GRCh38, and *Thalassospira lucentensis* Mv1 genomes.

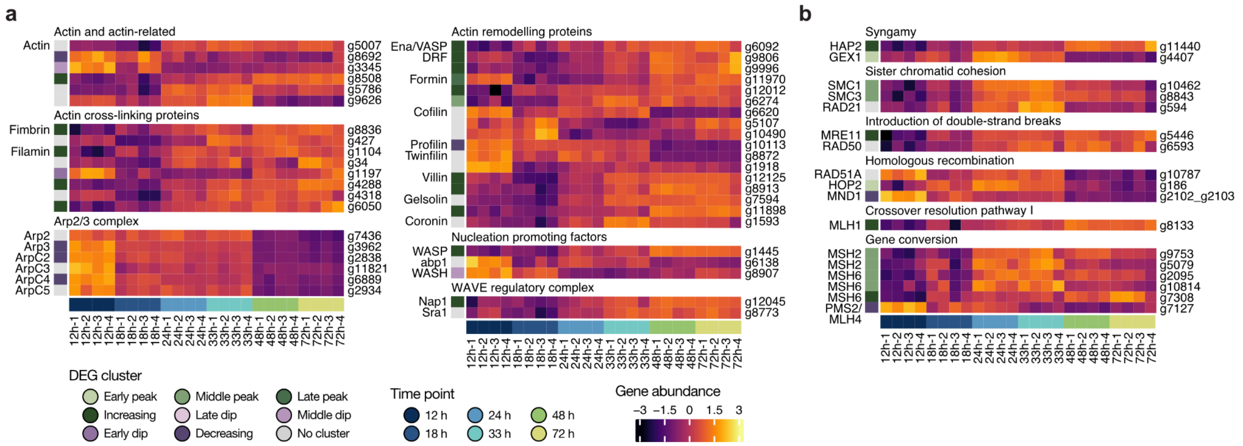


**Figure S5.** Heatmaps showing the expression patterns of genes involved in a) phagocytosis and b) meiosis that were significantly differentially expressed across the *M. vibrans* aggregation time series. The bottom row indicates the time point according to the legend and the left column DEG cluster assignment. Gene abundances for replicates at each time point were centered and scaled using the variance-stabilizing transformation of normalized gene counts.

**SUPPLEMENTARY DATA DESCRIPTIONS**

**Data S1.** 16S rDNA amplicon sequencing of xenic *M. vibrans* ATCC 50519.

**Data S2.** 16S rDNA sequence from the isolate *T. lucentensis* Mv1.

**Data S3.** 16S rDNA amplicon sequencing of monoxenic *M. vibrans* with *T .lucentensis* Mv1 generated from the ATCC 50519 strain.

**Data S4.** 16S rDNA amplicon sequencing of monoxenic *M. vibrans* with *T .lucentensis* Mv1 generated from the DSM 114853 strain.

**Data S5.** Summary of all sequence data submitted to NCBI under BioProject PRJNA1229532 with sample information, and BioSample and SRA accessions.

**Data S6.** Functional annotations of *M. vibrans* ATCC 50519 genes.

**Data S7.** RNA-sequencing gene expression counts including raw, normalized, and normalized and variance-stabilizing transformed data.

**Data S8.** Differentially expressed genes (DEGs) during *M. vibrans* ATCC 50519, including adjusted p-values, log2 fold change between 12 and 72 h, DEG cluster assignment, and functional annotations.

**Data S9.** Gene Ontology (GO) terms enriched in each DEG cluster.

**Data S10.** Multicellularity-related genes encoded in *M. vibrans* ATCC 50519, alongside annotation and predicted gene age information.

**Data S11.** Phagocytosis and meiosis-related genes encoded in *M. vibrans* ATCC 50519, alongside annotation information.

#### SUPPLEMENTARY VIDEO DESCRIPTIONS

**Video S1.** Aggregation of monoxenic *M. vibrans* ATCC 50519 with *T. lucentensis* Mv1 over 72h.

**Video S2.** Aggregation of aphidicolin-treated monoxenic *M. vibrans* ATCC 50519 with *T. lucentensis* Mv1 over 36h.

**Video S3.** Aggregation of monoxenic *M.vibrans* ATCC 50519 with *T. lucentensis* over 1140h (60 days).

**Video S4**. Aggregation of monoxenic *M.vibrans* DSM 114853 with bacterium *T. lucentensis* Mv1 over 72h.
